# Manganese availability determines insulin sensitivity by enhancing Akt activity

**DOI:** 10.64898/2026.09.02.746017

**Authors:** Jennifer R. Gamarra, Sei Higuchi, Timothy L. Yuan, Yuke Xie, Cong Liu, Niroshan Shanmugarajah, Hang Yang, Meredith O. Kelly, Sarah A. Hannou, Kathrin Schilling, Ana Navas-Acien, Eunhee Choi, Anum Glasgow, Inna I. Astapova, Mark A. Herman, Rebecca A. Haeusler

**Author notes:** Corresponding Author Information, Address: 1150 St Nicholas Ave, 6^th^ floor, New York, NY 10032. DECLARATION OF INTERESTS: The authors have declared that no conflict of interest exists.

## Abstract

Insulin signaling is a critical determinant of metabolic health, and impairments in insulin action contribute to the development of type 2 diabetes. The kinase Akt is a central mediator of insulin signaling and is required for insulin’s suppression of hepatic glucose output. Although the regulation of Akt by the insulin receptor-PI3K pathway is well understood, there are instances in which signaling downstream of Akt is dissociated from proximal insulin signaling, for example in insulin resistance. Nonetheless, little is known about PI3K-independent mechanisms of Akt regulation. Here, we discovered that hepatocyte manganese concentrations are a key determinant of PI3K-independent Akt function in vivo. We further demonstrated that manganese increases Akt’s catalytic efficiency, and quantitative phosphoproteomics revealed that manganese and insulin act additively to enhance Akt activity. Moreover, we uncovered that hepatic manganese concentrations fluctuate during fasting and feeding via carbohydrate-dependent transcriptional regulation of the manganese efflux transporter Slc30a10. This dynamic metal–signaling axis provides a mechanistic link between nutrient status and Akt activation, and suggests a molecular explanation for the glucose-lowering effects of manganese observed in humans. Our findings establish manganese as a physiologically regulated cofactor for Akt and position metal bioavailability as a previously unrecognized layer of insulin signaling control.

## INTRODUCTION

Nearly 50% of adults in the United States have diabetes or prediabetes (1), and insulin resistance is a defining pathologic feature of these conditions. At the molecular level, insulin resistance is predicted to be explained by defects in the distal elements of the insulin signaling pathway (2). Thus understanding insulin signaling–and the modulators of both the proximal and distal elements of the pathway–is fundamental to understanding diabetes pathogenesis.

The serine (Ser)/threonine (Thr)-protein kinase Akt is the essential mediator of distal insulin action. Insulin activates Akt by promoting its phosphorylation on two residues: Thr308 in the catalytic domain and Ser473 in the C-terminal tail (3, 4). These two modifications and the binding of Akt’s N-terminal pleckstrin homology (PH) domain to the membrane lipid phosphatidylinositol-3,4,5-triphosphate drive Akt from the autoinhibited to active state (5, 6). Phosphorylation of Thr308 and Ser473 are required for insulin-stimulated Akt activity (7), though other covalent post-translational modifications on Akt further tune activity (8). The activation of Akt by insulin results in decreased hepatic glucose output, which is impaired in insulin resistance and diabetes.

Kinases require divalent cations for enzymatic activity. A canonical function for divalent cations in kinases is to compensate for the negatively charged ATP used for phosphoryl transfer. The cation presumed to be involved in most kinase activity in vivo is magnesium (Mg^2+^), due to its high abundance and superior ability to support catalytic activity of model kinases including PKA (9). However, other divalent metals – such as manganese (Mn^2+^), cobalt, and cadmium – have been shown to support kinase activity in cell-free assays (9). Yet, the metalloproteome is not solved, and it remains uncertain which metals support the activity of which kinases in vivo.

Mn is essential for life and acquired through the diet. Cellular Mn concentrations are determined by the rate of cellular efflux, and the major mammalian Mn efflux transporter is Slc30a10 (10). Slc30a10 is present on the canalicular membranes of hepatocytes, where it mediates Mn efflux into the bile, and on the apical membranes of enterocytes, where it mediates Mn efflux into the intestinal lumen (11–13). It is also present in the brain where it protects against neurotoxicity (12), and rare congenital loss of *SLC30A10* function in humans causes neurologic deficits (14, 15). It is established that transcriptional regulation of *Slc30a10* determines cellular Mn efflux. Known factors that induce *Slc30a10* include hypoxia-inducible factors (16) and vitamin D receptor signaling in intestine (17–19). But despite its importance in biology, the regulation of Slc30a10 has been understudied.

In this study, we found that Mn acts on Akt to increase its activity in vivo, ex vivo, and in vitro, and this is sufficient to suppress glucose production. Furthermore, we found that *Slc30a10* expression and hepatobiliary Mn excretion is induced by dietary carbohydrates, due to direct binding of the carbohydrate response element binding protein (ChREBP) to the *Slc30a10* locus. These findings suggest that metal selectivity is a previously unexplored determinant of Akt activity that is regulated by carbohydrate metabolism.

## RESULTS

### Slc30a10^Tbg,Vil^ mice have increased liver Mn and low blood glucose without increased insulin

We used mice with floxed alleles of the Mn efflux transporter Slc30a10 (13). Previous research established that Slc30a10-mediated transintestinal excretion of Mn can compensate for knockout of Slc30a10 in hepatocytes alone (13, 20). Therefore, to modulate hepatic Mn levels we knocked out Slc30a10 from intestinal epithelium (Villin-Cre) and hepatocytes (AAV8-Tbg-Cre) to cause specific loss of *Slc30a10* in liver and intestine (**Supplemental Figure 1A**) (hereafter, Slc30a10^Tbg,Vil^ mice). Controls were sex-matched littermate Slc30a10^fl/fl^ mice transduced with AAV8-GFP. Slc30a10^Tbg,Vil^ mice showed increased concentrations of Mn in liver and blood (**Figure 1A, Supplemental Figure 1B**). Soleus muscle from Slc30a10^Tbg,Vil^ mice also showed increased Mn compared to controls, however the Mn concentrations in soleus and epididymal white adipose tissues were substantially lower than liver (**Supplemental Figure 1C**). Slc30a10^Tbg,Vil^ mice showed no differences in other liver metals except increased iron, reflecting increased red blood cell hemoglobin (21), and no differences between genotypes in any other metals (**Supplemental Figure 1D-I**). Though human congenital loss of SLC30A10 function is associated with cirrhosis (14, 15), Slc30a10^Tbg,Vil^ mice showed normal serum AST and ALT and other markers of liver function (**Supplemental Figure 1J-O)**, indicating there is no overt liver injury. There was a significant increase in red blood cells in Slc30a10^Tbg,Vil^ mice (**Supplemental Figure 1P**), consistent with polycythemia observed in people with *SLC30A10* loss of function variants (14, 15).

**Figure 1.**
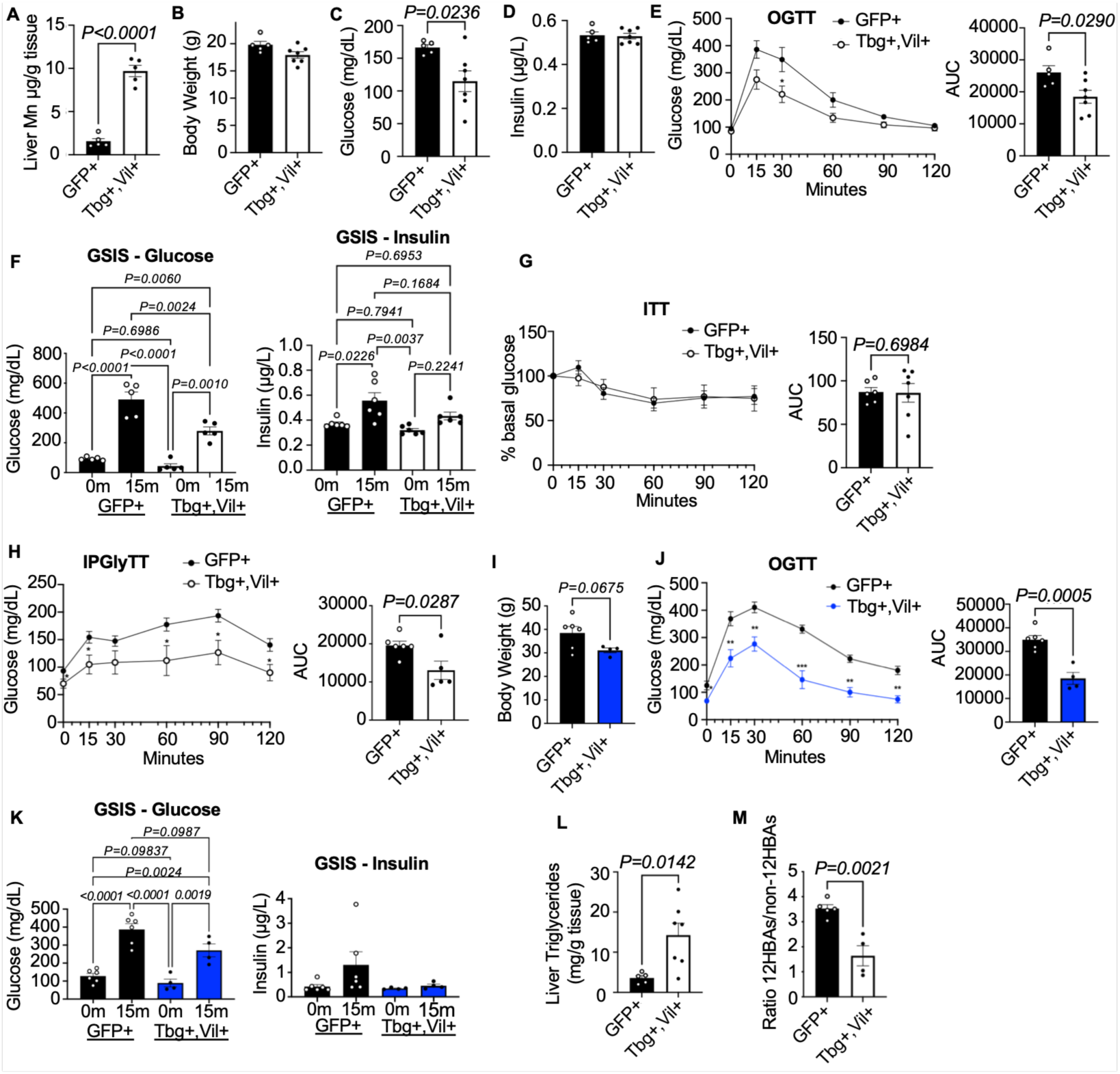
Metabolic parameters in Slc30a10^Tbg,Vil^ mice. (A) Liver Mn, n=5 males/group. (B) Body weight, 5-hour fasting (C) blood glucose, (D) plasma insulin, and (E) oral glucose tolerance test (OGTT), n=5-6 females/group. (F) Blood glucose and plasma insulin during glucose-stimulated insulin secretion (GSIS) tests, (G) insulin tolerance test (ITT), (H) intraperitoneal glycerol tolerance test (IPGlyTT), n=5-6 males/group. (I) Body weight, (J) OGTT, (K) GSIS in mice fed a high-fat, high-sucrose diet for 14 weeks, n=4-6 males/group. (L) Liver triglycerides and (M) ratio of 12α-hydroxylated bile acids (12HBAs) to non-12HBAs in the total bile acid pool of chow diet fed mice, n=5-7 males/group. Data presented as mean ± SEM. ns=not significant, *p<0.05, **p<0.01, ***p<0.001, ****p<0.0001 by Student’s t-test except (F) and (K) where two-way ANOVAs were used.

There were no significant differences in body weight between Slc30a10^Tbg,Vil^ and controls (**Figure 1B, Supplemental Figure 2A**). After a 5-hour fast Slc30a10^Tbg,Vil^ mice had significantly lower glucose levels, but there were no differences in plasma insulin or liver glycogen (**Figure 1C-D, Supplemental Figure 2B**). After oral glucose gavage Slc30a10^Tbg,Vil^ female and male mice showed lower blood glucose than controls (**Figure 1E, Supplemental Figure 2C**). We observed similar results after intraperitoneal glucose injection (**Supplemental Figure 2D**). After pooling data from multiple cohorts, we confirmed that there are no differences in body weight between genotypes (20.6 ± 0.82 vs 22.1 ± 0.49 for Slc30a10^Tbg,Vil^ mice vs controls, n = 30-32, P=0.13), and low body weight does not explain the reduced glycemia in Slc30a10^Tbg,Vil^ mice (**Supplemental Figure 2E-F**).

We next examined glucose-stimulated insulin secretion. 15-minutes after oral glucose, Slc30a10^Tbg,Vil^ mice had significantly lower glucose than controls, but there were no differences in plasma insulin appearance or free fatty acid suppression between groups (**Figure 1F, Supplemental Figure 2G**). In insulin tolerance tests – which primarily reflect muscle and adipose glucose uptake – we observed no differences between groups (**Figure 1G**). In glycerol tolerance tests – which assess gluconeogenesis – Slc30a10^Tbg,Vil^ mice showed lower blood glucose throughout the test, suggesting decreased gluconeogenesis (**Figure 1H**).

We next fed mice a high-fat, high-sucrose diet. There were no differences in body weight, 5-hour fasting blood glucose or insulin between groups (**Figure 1I, Supplemental Figure 2H-I**). However, Slc30a10^Tbg,Vil^ mice showed significantly lower glucose after oral or intraperitoneal glucose bolus, compared to controls, without increased insulin secretion (**Figure 1J-K, Supplemental Figure 2J**). We found no significant differences in glucose tolerance in Slc30a10^Tbg^ or Slc30a10^Vil^ mice compared to controls fed a standard chow diet or a high fat, high sucrose diet (**Supplemental Figure 3A-D**). Overall, these results indicate that hepatic Mn accumulation results in lower glycemia without increased insulin. This phenotype is similar to what is observed in mouse models of increased insulin action (22–24) and is an established indicator of insulin sensitivity.

### Slc30a10^Tbg,Vil^ mice show multiple features of increased hepatic insulin action

A potential explanation for reduced glycemia is increased activation of hepatic insulin signaling. To examine this, we assessed other metabolic markers of increased hepatic insulin action. Insulin signaling promotes liver triglyceride accumulation, partly by Akt-dependent induction of lipogenesis (25). Consistent with this, Slc30a10^Tbg,Vil^ mice showed high liver triglycerides, but no differences in other lipid parameters (**Figure 1L, Supplemental Figure 3E-G**). Hepatic insulin signaling also suppresses the subset of bile acids that are hydroxylated at carbon 12α (26–28). Consistent with this, Slc30a10^Tbg,Vil^ mice showed lower 12α-hydroxylated bile acids but normal total bile acid pool size (**Figure 1M, Supplemental Table 1**).

### Mn increases hepatic insulin signaling

To directly determine whether Slc30a10-depletion enhances the insulin signaling cascade, we injected mice with insulin via the jugular vein and collected insulin sensitive tissues including liver, adipose and muscle. In liver, there were no differences in insulin receptor protein expression or phosphorylation between genotypes (**Figure 2A-B**). There were also no differences in Akt protein expression, though livers of Slc30a10^Tbg,Vil^ mice had higher phosphorylation at Thr308 (**Figure 2A-B**). A notable difference between genotypes was an increase in the phosphorylation of Akt substrates, including defined targets GSK3β and PRAS40 (**Figure 2A-B**), and diverse Akt target proteins, as detected by an antibody against the phosphorylated Akt consensus motif (**Figure 2C-D**). In contrast, in epididymal white adipose tissue and soleus muscle there were no differences in phosphorylation of Akt between groups (**Supplemental Figure 4A-B**). There was a significant increase in GSK3β phosphorylation in soleus muscle of Slc30a10^Tbg,Vil^ mice (**Supplemental Figure 4A-B)**, consistent with an increase in soleus Mn levels (**Supplemental Figure 1C)**. There were no differences in the liver activity of the highly related kinase protein kinase A (PKA) (**Supplemental Figure 4C-D**). These results demonstrate that mice with increased liver Mn have increased activation of the Akt signaling pathway.

**Figure 2.**
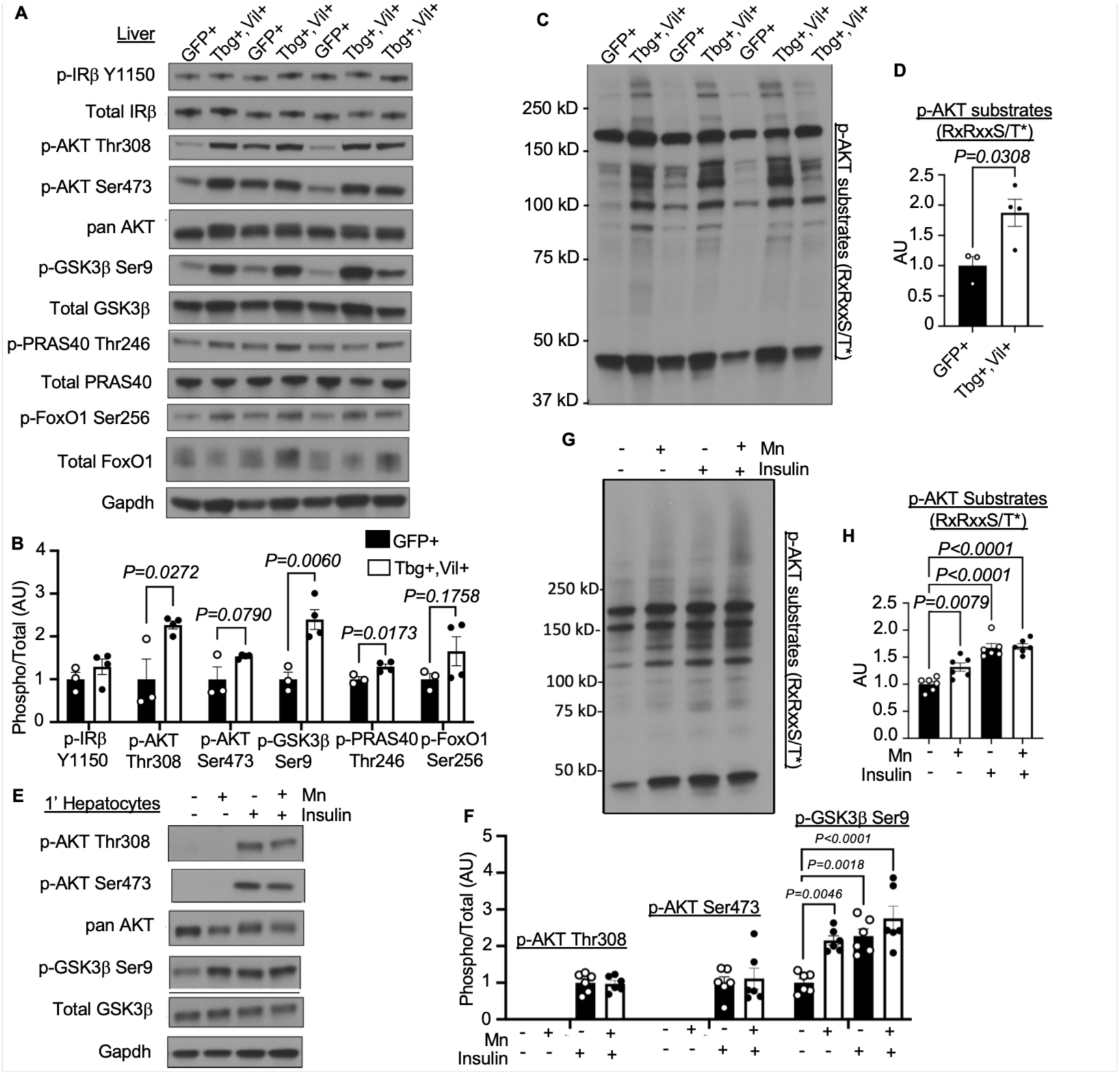
Slc30a10^Tbg,Vil^ livers and Mn-treated hepatocytes have increased Akt signaling. Immunoblots and densitometric quantification of (A-B) the insulin signaling pathway and (C-D) phosphorylated Akt substrates in livers from insulin-injected Slc30a10^Tbg,Vil^ mice. Immunoblots and densitometric quantification of (E-F) the insulin signaling pathway and (G-H) phosphorylated Akt substrates in primary mouse hepatocytes treated with Mn (6 μM) or insulin (1 nM). (E-H) Display representative blots from three independent experiments. Data presented as mean ± SEM. Student’s t-test for (A-D) and one-way ANOVA for (E-H) were used.

We next tested whether Mn increases Akt signaling in a hepatocyte-autonomous manner using primary murine hepatocytes. Mn treatment in hepatocytes led to a specific increase in intracellular Mn, without changes in Fe (**Supplemental Figure 4E-F**). As expected, insulin stimulated the phosphorylation of Akt at canonical sites Thr308 and Ser473 and phosphorylation of Akt substrates (**Figure 2E-H**). We observed that Mn treatment was also sufficient to stimulate Akt activity - as evidenced by significant increases in phosphorylation of Akt substrates - without appreciable phosphorylation of Akt at Thr308 or Ser473 (**Figure 2E-H**). There were no differences in PKA activity (**Supplemental Figure 4G**).

To determine whether observed increases in Akt activity had functional consequences on cellular metabolism, we measured glucose production in primary hepatocytes. As expected, glucose production was induced by forskolin/dexamethasone and suppressed by insulin (**Figure 3A**). We found that Mn treatment alone was sufficient to suppress glucose production (**Figure 3A**).

**Figure 3.**
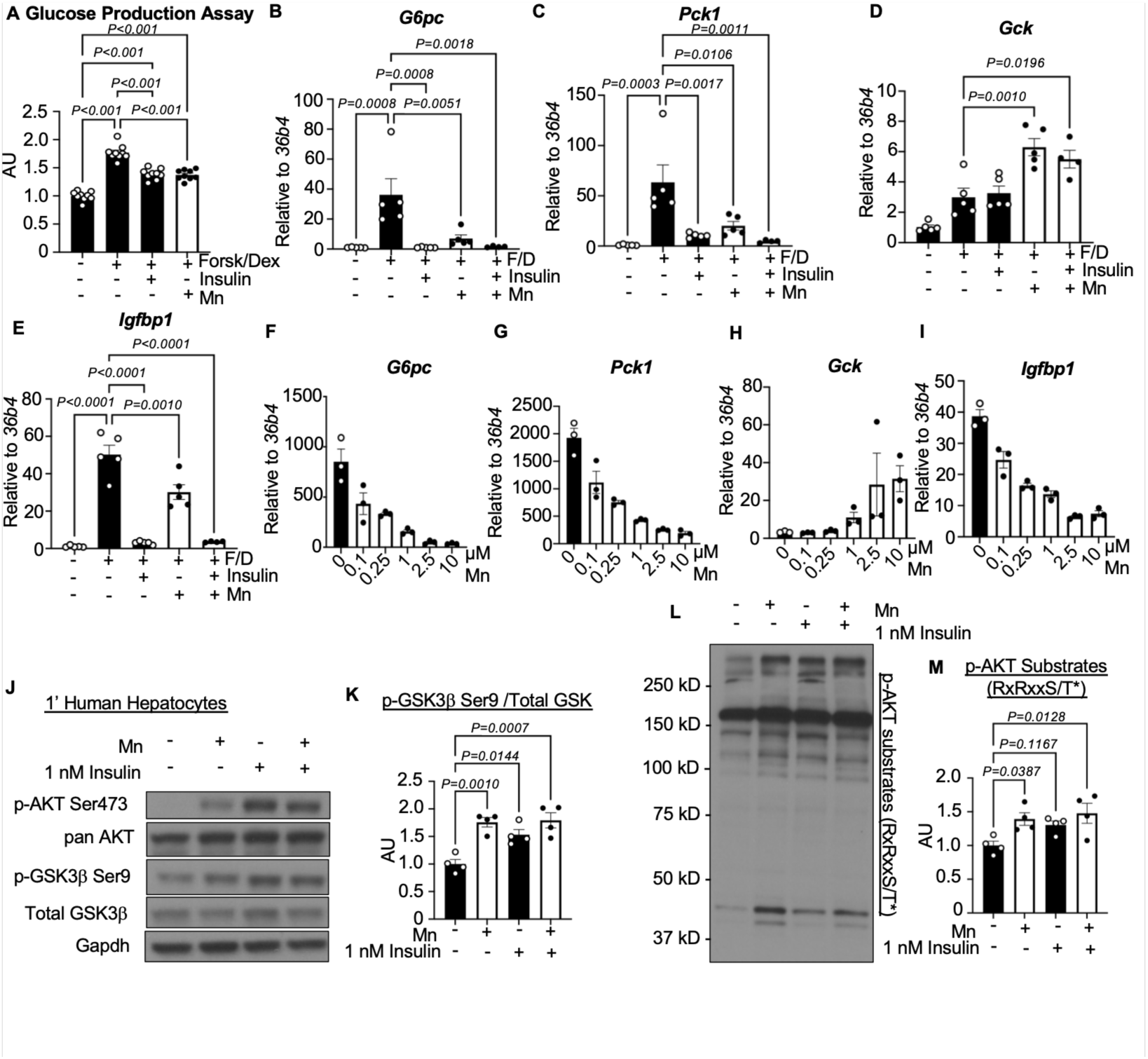
Mn-treated hepatocytes have increased Akt signaling with decreased glucose production and gluconeogenic gene expression. (A) Glucose production and (B-I) gene expression from primary hepatocytes treated with forskolin/dexamethasone (Forsk/Dex), Mn (6 μM) or insulin (1 nM). (J-M) Representative immunoblots from primary human hepatocytes treated with Mn (6 μM) or insulin (1 nM) with relative densitometric quantifications. Each data point represents a technical replicate from the same donor. Data presented as mean ± SEM. One-way ANOVAs were used.

Akt suppresses glucose production cell autonomously in hepatocytes by inactivation of FoxO transcription factors, thus downregulating glucose-6-phosphatase catalytic subunit (*G6pc*), phosphoenolpyruvate carboxykinase 1 (*Pck1*), and insulin like growth factor binding protein 1 (*Igfbp1*), and upregulating glucokinase (*Gck*). We observed that Mn was sufficient to mimic the effects of insulin on these transcripts (**Figure 3B-E**), and many other insulin-regulated transcripts and pathways (**Supplemental Figure 4H, Supplemental Tables 2-3**). The effects on glucoregulatory genes occurred dose-dependently in response to physiologically-relevant Mn concentrations (**Figure 3F-I**). The effect of Mn to promote phosphorylation of Akt targets was reproducible in independent experiments in murine primary hepatocytes (**Supplemental Figure 5A-G**) and human primary hepatocytes repeated with multiple donors (**Figure 3J-M, Supplemental Figure 5H-J**). These findings indicate that in vivo in livers and ex vivo in primary hepatocytes, Mn increases Akt activity and suppresses glucose production.

### Mn acts on Akt to increase kinase activity

To confirm that the effects of Mn to promote phosphorylation of Akt targets occurs via Akt specifically, we used the ATP-competitive Akt inhibitor ipatasertib. Ipatasertib was sufficient to block the effects of both insulin and Mn on Akt target phosphorylation (**Figure 4A-B, Supplemental Figure 6A**). We also reproduced the known paradoxical effect of ipatasertib to promote Akt phosphorylation (29, 30) (**Figure 4A**). These results indicate that the effect of Mn to induce phosphorylation of Akt targets requires Akt activity.

**Figure 4.**
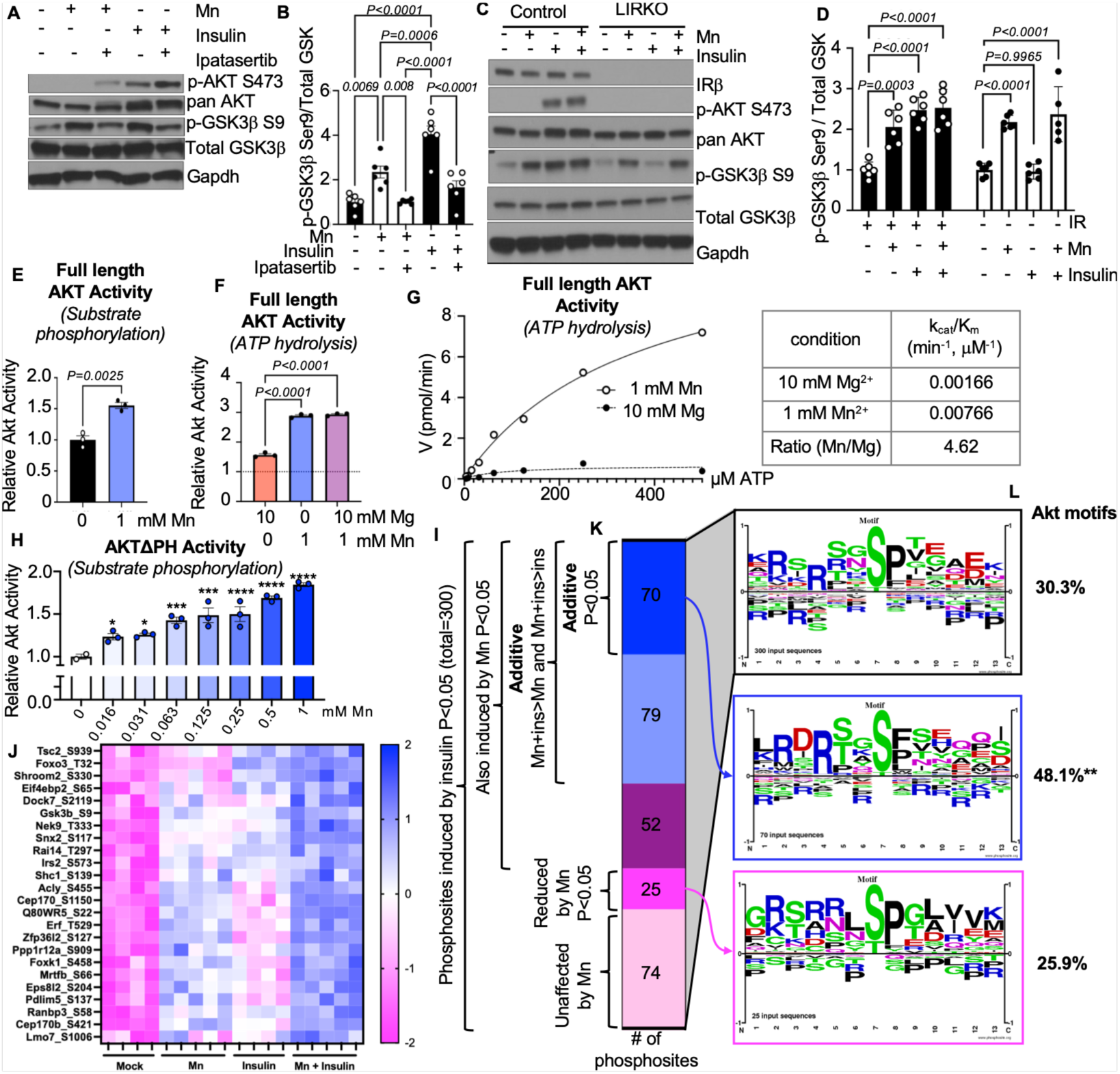
Mn increases Akt catalytic activity and is additive with insulin. Representative immunoblots of the insulin signaling pathway from (A) primary hepatocytes treated with insulin (1 nM), Mn (6 μM), or ipatasertib (10 μM) with (B) densitometric quantification, and (C) from primary hepatocytes derived from control or LIRKO mice treated with insulin or Mn with (D) densitometric quantification. (E-G) In vitro activity of full-length Akt. Enzyme catalytic efficiency in (G) is reported as k_cat_/K_m_ and is representative of three independent experiments. (H) In vitro activity of AktΔPH. (I-L) Phosphoproteomic analysis in murine primary hepatocytes. For details, see “Phosphoproteomics data analysis” in the methods section. (I) Phosphosites induced by insulin (1 nM) in mouse primary hepatocytes (p<0.05), subdivided by the effect of Mn (6 μM). (J) Heatmap of select significantly additive phosphosites. (K) Sequence logos from peptides in the indicated category. (L) Percent of phosphopeptides in the indicated category containing an Akt recognition motif (RxRxxS/Tϕ, RxRxxS/T, RxxS/Tϕ, or RxxS/T; ϕ=hydrophobic residue). Data presented as mean ± SEM. Student’s t-test for (E), one-way ANOVA for (B, D, F, H), moderated t-test (75) for (I) and Treat method(37) for additivity in (I, J), and Fisher’s exact test for (L) were used. *p<0.05, **p<0.01, ***p<0.001, ****p<0.0001.

One possible explanation for Mn-induced Akt activity is increased upstream signaling through the insulin receptor axis. To directly test the role of upstream insulin signaling, we used insulin receptor-floxed mice transduced with AAV8-Tbg-Cre (LIRKO). In primary hepatocytes from LIRKO mice, insulin receptor was absent, and insulin treatment failed to phosphorylate Akt or its target GSK3β (**Figure 4C-D**). Yet, even in the absence of insulin receptor or appreciable Akt phosphorylation, Mn was still able to increase phosphorylation of GSK3β (**Figure 4C-D**). Moreover, while insulin’s ability to suppress *G6pc* and *Pck1* was impaired in LIRKO primary hepatocytes, Mn still fully suppressed those transcripts (**Supplemental Figure 6C-D**). These results indicate that Mn is unlikely to function upstream of Akt.

A second potential explanation is that Mn acts directly on Akt to increase its activity, and MAHOMES II (31) predicts Mn binding at catalytic sites on Akt. AGC family kinases contain two conserved metal binding residues in the kinase domain, which are present in Akt: Asn279 in the catalytic loop and Asp292 in the activation loop (residue numbering for human AKT1). Metals bound at these sites counteract the negative charges of ATP, aid in the positioning of the phosphate and substrate for phosphoryltransfer, and residue Asp292 is required for kinase activity (32). Indeed, Mn ions are found bound to both Asn279 and Asp292 in crystal structures of Akt (30, 33).

To address this, we tested the direct effects of Mn on the activity of purified, full-length Akt. We found that, in combination with ATP, Mn was sufficient to induce the activity of full-length Akt in vitro to phosphorylate GSK3β and to consume ATP, even in the presence of tenfold excess Mg (**Figure 4E-F, Supplemental Figure 6E**). This was due to a ∼4.6-fold increase in the catalytic efficiency (k_cat_/K_m_) of Akt in the presence of Mn compared to Mg (**Figure 4G, Supplemental Figure 6F**). This effect was not universal across kinases, as PKA was preferentially activated by Mg compared to Mn (**Supplemental Figure 6G**), consistent with our results in mice and primary hepatocytes.

Full-length Akt is auto-inhibited by its lipid-binding PH domain, which limits substrate access to the catalytic domain. Thus we assessed the effects of Mn on the activity of purified Akt lacking the PH domain (AktΔPH). We observed that micromolar concentrations of Mn were sufficient to activate AktΔPH (**Figure 4H**). Altogether, these results demonstrate that Mn acts directly on Akt protein to simulate its activity by increasing the catalytic efficiency.

### Mn and insulin act additively on Akt

In vivo, there are multiple extrahepatic inputs (34–36) to ensure hepatic insulin signaling is inactive during fasting and then robustly activated by insulin. Our experiments in mice demonstrate that insulin-stimulated Akt activity is further activated in an additive manner by elevations in liver Mn (see Figure 2A).

To quantitatively assess (a) whether insulin and Mn act additively on the phosphorylation of Akt targets in primary hepatocytes, and (b) gain insight into whether Mn affects the specificity of proteins phosphorylated by the insulin signaling cascade, we performed phosphoproteomics. We found that insulin significantly induced 300 unique phosphosites (**Figure 4I, Supplemental Table 4**). Of those, 149 (∼50%) phosphosites were also significantly induced by Mn *and* the effect size of Mn + insulin was greater than the effect size of either treatment alone, and we considered those sites to be additively phosphorylated (**Figure 4I**). We statistically analyzed the phosphosites considering the effect size of additivity using the Treat method (37), which identified the 70 top additive phosphosites (**Figure 4I**). These additive sites included many established targets of hepatic insulin signaling (38) (**Figure 4J, Supplemental Table 4**).

We analyzed the phosphopeptide sequences and found that insulin-induced phosphosites were strongly enriched for motifs recognized by Akt, including the canonical RxRxxS/T* (**Figure 4K-L, Supplemental Table 4**). Additive phosphosites showed overrepresentation of Akt motifs and a preference for Asp in the −4 position (**Figure 4K-L, Supplemental Table 5**). We also performed pathway analysis for kinase and phosphatase substrates and found significant increases in activity of kinases involved in insulin signaling including AKT, mTOR and PDK1, and a decrease in activity in phosphatases that inhibit insulin signaling such as PTEN within the additive sites (**Supplemental Table 6**). Altogether, this analysis indicates that (a) Mn acts additively with insulin to promote the phosphophorylation of peptide substrates and (b) the additively phosphorylated peptides are enriched for Akt recognition motifs.

### ChREBP-dependent carbohydrate signaling induces Slc30a10 and hepatobiliary Mn efflux

A crucial physiologic setting when hepatic Akt must be robustly activated is in the transition from fasting to feeding. Postprandial insulin induces signaling through the hepatic insulin receptor-Akt cascade within seconds to minutes (38). Thus, at the time of the transition between fasting and feeding, the components of the insulin signaling cascade, including Akt, should be primed for maximal activation potential. Therefore, we investigated whether hepatic Mn availability is regulated by fasting and feeding. We fasted wild-type chow-fed mice for up to 24-hours, or refed them following a 24-hour fast. We observed low hepatic *Slc30a10* expression after fasting and a robust induction upon refeeding, with highest expression occurring after 4-hours (**Figure 5A**). Liver Mn importers *Slc39a8* and *Slc39a14* were unchanged during the fasting-refeeding transition (**Figure 5A**). We observed similar effects in mice fed a western-type diet (**Supplemental Figure 7A**). We tested the functional consequences of the nutritional regulation of *Slc30a10*. We found that hepatic Mn concentrations were high in fasting, aligning with high glucose and insulin levels, and – consistent with induction of the efflux transporter – decreased hours after refeeding (**Figure 5B, Supplemental Figure 7B-C**). These findings indicate that after fasting, Mn accumulates in liver, whereas refeeding causes induction of *Slc30a10* and Mn egress into the bile, in a process that proceeds over multiple hours.

**Figure 5.**
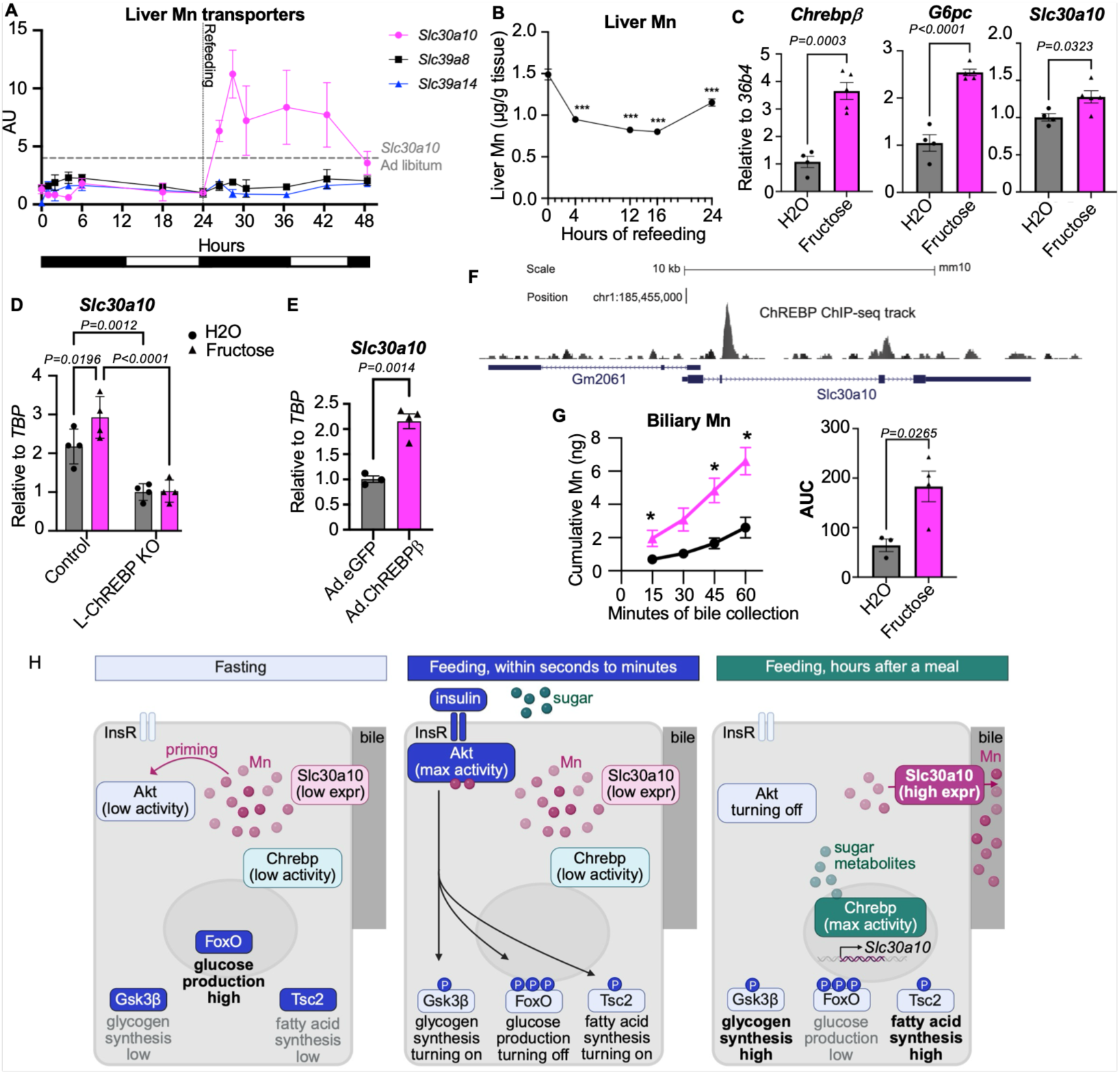
*Slc30a10* and hepatobiliary Mn efflux is nutritionally regulated by ChREBP. (A) Liver expression of Mn transporters in C57BL/6J mice fasted for 24-hours followed by refeeding with chow diet, n=3-5 males/group. Each gene is plotted relative to its expression at the 24h-fast. Dashed line denotes *ad libitum Slc30a10* expression. (B) Liver Mn concentrations during refeeding with chow diet after a 24-hour fast in C57/BL6J mice, n=5 males/group. Liver gene expression from (C) wild type C3H/HeJ mice gavaged with fructose or water, n=4-5 males/group, (D) control and liver-specific ChREBP knockout mice (L-ChREBP KO) mice gavaged with fructose or water, n=4 females/group, and (E) C57BL/6J mice transduced with indicated adenoviruses, by quantitative PCR, n=3-4 males/group. (F) ChREBP ChIP-Seq signal track near *Slc30a10* gene in liver of male C3H/HeJ mice 90 minutes after fructose gavage. (G) Cumulative biliary Mn in C3H/HeJ mice 2-hours after fructose or water gavage, n=3-4 males/group. (H) Proposed model of Mn regulation of Akt activity and hepatobiliary Mn efflux by ChREBP made in BioRender. AUC = area under the curve. Data are presented as mean ± SEM. Student’s t-tests for all panels except (D) by one-way ANOVA were used. In panel (B) a Student’s t-test was performed relative to time 0. In panel (G) a Student’s t-test was performed against fructose gavage at each time point. *p<0.05, **p<0.01, ***p<0.001, ****p<0.0001.

We next considered which signals of fasting and feeding were responsible for regulating *Slc30a10*. We reasoned that the transcription factor carbohydrate-responsive element-binding protein (ChREBP) is a candidate to explain the transcriptional regulation of *Slc30a10*. Its activity is low during fasting and highly induced by carbohydrates. To test the role of ChREBP, we performed fructose gavage in wild-type C3H/HeJ mice, which is known to acutely activate ChREBP (39). Fructose gavage caused the expected upregulation of *Chrebpβ* and *G6pc* (39) in liver, as well as upregulation of *Slc30a10*, which required ChREBP (**Figure 5C-D**). Moreover, overexpression of ChREBPβ induced *Slc30a10* expression in liver and primary hepatocytes (**Figure 5E, Supplemental Figure 7D**). Finally, chromatin immunoprecipitation showed that ChREBP directly binds the *Slc30a10* genomic locus (**Figure 5F**). The binding site is downstream of the transcription start site, in the first intron, which is a common binding region for ChREBP and other transcription factors (40–43).

We next tested the functional consequences of fructose-dependent induction of *Slc30a10*. We found that fructose gavage in C3H/HeJ mice was sufficient to triple the rate of biliary Mn excretion, with no differences in bile flow rate (**Figure 5G, Supplemental Figure 7E**). There were no effects of fructose on the biliary efflux of copper (**Supplemental Figure 7F**), another metal that is excreted primarily through the hepatobiliary route (44). Collectively, these data indicate that ChREBP directly induces *Slc30a10* and promotes hepatobiliary Mn excretion.

## DISCUSSION

Here we demonstrate that hepatic Akt activity is induced by Mn in vivo, ex vivo, and in vitro. Moreover, we demonstrate that hepatobiliary Mn efflux, which is a principal determinant of Mn homeostasis, is induced by carbohydrate signaling via direct regulation of *Slc30a10* by ChREBP, in a process that proceeds over multiple hours. One might speculate that the delayed reduction of liver Mn after a meal may contribute to the termination of insulin signaling and the prevention of hypoglycemia. Overall, these data suggest that controlled access to Mn^2+^ ions is a novel, nutritionally-controlled layer of hepatic Akt regulation that is active in vivo. A model is shown in **Figure 5H**.

Mn is a trace element known for its role as a cofactor for a variety of enzymes. Among the best-known Mn-dependent enzymes are arginase, the manganese-dependent superoxide dismutase, and enzymes involved in glycosylation (45, 46). It is well-established that Mn is essential and must be maintained at minimum levels, yet excess Mn is detrimental, and the latter is exemplified by rare, congenital loss-of-function mutations in *SLC30A10* (14, 15). While recent research has greatly advanced our knowledge of Mn transport (47–49), our understanding of the physiological effects of Mn has lagged behind. This work partially advances us towards the goal of understanding physiological roles of Mn.

Although Slc30a10^Tbg,Vil^ mice do not model a human disease, they are a tool to investigate the effects of Mn. Using this tool, we have now been able to provide a molecular mechanism for prior observations that Mn has glucose-lowering effects in humans and mice. The first medical report that Mn lowers blood glucose was published in the *Lancet* in 1962 (50). This report inspired a study in 1966 demonstrating that male patients with chronic Mn ore exposure (n=11) had significant hypoglycemia in intravenous glucose tolerance tests, compared to healthy controls (n=20) (51). More recently, it was reported that in mice fed high-fat diet, 8 weeks of 5x per week intraperitoneal injection with MnCl_2_ substantially improved glucose tolerance (52). These reports, together with our findings, establish that elevating liver Mn by several independent mechanisms in humans and mice has reproducible glucose-lowering consequences.

Due to high concentrations of Mg^2+^ in cells (53), it is widely believed that Mg-ATP is the nucleotide complex used in most kinases in cells and organisms. Indeed, Mg^2+^ supports the highest activity of PKA, compared to Mn^2+^ and other metals (9, 54). Yet this dogma may not hold true for every kinase. Intriguingly, two kinases that escape the dogma are upstream and downstream of Akt: for both the insulin receptor itself and target of rapamycin complex 1 (TORC1), experiments in cell-free systems have shown that Mn^2+^ activates their kinase activity (55–57). These findings are supported by experiments in cell lines and yeast (57–59). Taken together with our finding that Mn improves the catalytic efficiency of Akt, this suggests the possibility that Mn bioavailability supports the coordinated activation of multiple kinases in the insulin signaling cascade. It will be of interest to identify additional Mn-sensitive kinases in future investigations.

Our findings indicate that hepatic Mn is dynamically controlled during the fasting-feeding transition. In normal physiology, this pathway is mediated by carbohydrate-induced ChREBP signaling. Notably, a recent study described transcriptional activation of *Slc30a10* by chronic high-fructose diet, in a ChREBP-dependent manner (60). Our work extends this observation by showing that (i) this pathway is sufficient to stimulate biliary Mn efflux, (ii) the induction of *Slc30a10* occurs in response to other forms of dietary carbohydrates–as chow and western diets contain sucrose and starch, but not high doses of pure fructose, and (iii) the induction of *Slc30a10* and biliary Mn efflux occurs within hours, not just in response to chronic excess fructose.

Overall, our work has identified a distinct role for Mn in Akt signaling. In future investigations, it will be interesting to determine the effects of this regulatory axis, and its potential modulation, on diabetes and other metabolic diseases. Our work also highlights an underappreciated role for metal ion selectivity in kinase function in vivo. This may lead to novel implications for basic research, kinase-directed therapeutics, and metal ion-directed therapeutics.

## METHODS

### Sex as a biological variable

For experiments with Slc30a10^fl/fl^ mice both male and female sexes were used and data are reported from both. For experiments with fructose and ChREBP modulation a mixture of male and female sexes were used and figure legends denote the sex of the mice used in the data displayed.

### Mice and diets

Mice were housed in a facility with a 12-h light/12-h dark cycle and had free access to chow (3.4 kcal/g, Purina 5053; 24.7% kcal from protein, 62.1% from carbohydrate, and 13.2% fat). Slc30a10^fl/fl^ mice were kindly shared by Thomas Bartnikas (13). We crossed Slc30a10^fl/fl^ mice with mice bearing Villin-Cre. Mice were genotyped using a genotyping kit (KAPA Biosystems #KK5621). The DNA products after PCR were resolved by electrophoresis on a 2% agarose gel with 0.05% ethidium bromide. At 4 weeks of age, we injected sex-matched littermate mice intravenously with 10^11^ viral particles/mouse of AAV8-Tbg-Cre (Addgene #107787-AAV8) to Slc30a10^Vil^ mice or AAV8-Tbg-GFP (Addgene #105535-AAV8) to littermate Slc30a10^fl/fl^ mice as controls. Experiments began 3 weeks after AAV transduction. Generation and experiments using liver-specific ChREBP knock-out mice were described previously (61). Liver-specific insulin receptor knockout mice were generated using Insr^fl/fl^ that had been backcrossed 9 generations onto C57BL/6J and injecting them with 10^11^ viral particles/mouse of AAV8-Tbg-Cre or AAV8-Tbg-GFP. Hepatocytes were isolated two weeks after AAV transduction. Wild type C57BL/6J (#000664) and C3H/HeJ (#000659) mice were from Jackson Laboratories. For experiments of in vivo ChREBPβ overexpression, wild type C57BL/6J mice were injected retro-orbitally at a dose of 5 x 10^10^ viral particles/mouse with an adenovirus expressing eGFP (Ad.eGFP) or an adenovirus expressing 3xFlag-ChREBPβ. Mice were sacrificed and livers collected 8 days after injections Generation of Ad-ChREBPβ has been described previously (62). During high-fat, high-sucrose diet interventions, mice were weaned onto 60% kcal from fat diet (Research Diets #D12492) for 14 weeks. For fasting-refeeding experiments, mice were fed either chow or western diet (Envigo TD.88137) for one week prior to the experiment.

Mice were euthanized using CO_2_ asphyxiation followed by secondary cervical dislocation. Blood was collected through the inferior vena cava. Whole blood was aliquoted into metal-free tubes (Labcon North America # 31343450019), the remainder was put in EDTA-coated tubes to obtain plasma. Prior to collecting perfused tissues, mice were perfused via the heart with PBS. After perfusion, liver, epididymal white adipose tissue, and soleus muscle were excised into metal-free tubes.

### Fructose gavage

Mice were fasted for 5-hours prior to oral gavage with water or 4 g/kg body weight fructose. Mice were euthanized 100 minutes after gavage.

### Fasting-refeeding

Wild type C57BL/6J male mice were synchronized by removing all food during 10:00 – 17:00 hours, then chow diet or western diet was replenished at 17:00 hours. Fasting began at 19:00 hours. Mice were euthanized and liver tissue was collected at time points from 0 up to 24-hour fasting, or 24-hour refeeding.

### Glucose tolerance and glucose stimulated insulin secretion tests

Mice were fasted in fresh cages prior to being orally gavaged or intraperitoneally injected with 2 g/kg body weight glucose. Blood glucose measurements were recorded via tail vein bleeding using Contour Next One Blood Glucose Monitor and strips (Contour) at 0, 15, 30, 60, 90 and 120 minutes before replenishing diet. For insulin and non-esterified fatty acid measurements, blood was collected by submandibular bleeding into EDTA-coated tubes.

### Insulin tolerance tests

Mice were fasted for five hours in fresh cages prior to being intraperitoneally injected with 0.5 U/kg body weight (Novolog, Novo Nordisk). Glucose measurements were recorded via tail vain bleeding using OneTouch glucose monitor and strips (Contour) at 0, 15, 30, 60, 90 and 120 minutes.

### Glycerol tolerance tests

Mice were fasted in fresh cages prior to being intraperitoneally injected with 2 g/kg body weight glycerol in saline. Glucose measurements were recorded via tail vain bleeding using OneTouch glucose monitor and strips (Contour) at 0, 15, 30, 60, 90 and 120 minutes.

### In vivo insulin stimulation tests

Mice were fasted for 5-hours prior to anesthetizing using ketamine (100 mg/kg)/xylazine (10 mg/kg) IP injection. Mice were affixed and the jugular vein was exposed for injection of 2 U/kg body weight insulin (Novolog, Novo Nordisk) using an insulin syringe. After 15 minutes, the body cavity was opened and the liver was collected. At the 17 minute time point, the epididymal white adipose tissue was collected and at 19 minutes the soleus muscle was collected.

### Liver metabolic tests

Hepatic lipids were extracted as described (63) and measured using colorimetric assays for triglycerides (Infinity; Thermo Scientific #TR22421) and cholesterol (Cholesterol E, Wako Diagnostics # 99902601).

To measure hepatic glycogen content, 100 mg of liver tissue was used. 500 *μ*L of 6% perchloric acid was added and homogenized using a rotor-stator homogenizer. The sample was centrifuged at top speed for 10 mins at 4C. 250 *μ*L of the supernatant was transferred to a new vial with 250 *μ*L of deionized water. The sample was centrifuged at top speed for 10 mins at 4C. 400 *μ*L of the supernatant was transferred to a new vial where the pH was adjusted to between 6 – 7 using KOH. Sample was centrifuged at top speed for 10 mins at 4C before processing for glycogen breakdown by incubating 20 *μ*L of sample with 100 *μ*L of amyloglucosidase (1 mg/mL) for 2 hours shaking at 42C. The glucose released in the glycogen breakdown was quantified using Sigma’s Glucose Assay Kit (HK glucose assay, #GAHK-20-1KT).

### Plasma metabolic tests

Plasma lipids were measured using colorimetric assays for triglycerides (Infinity; Thermo Scientific #TR22421), cholesterol (Cholesterol E, Wako Diagnostics # 99902601), non-esterified fatty acids (NEFA) (Wako Diagnostics # 999-34691, #995-34791, # 991-34891, # 993-35191). Plasma insulin was measured using an ELISA kit (Mercodia, #10-1247-01). Plasma alanine aminotransferase (ALT) (Sigma, #MAK052), aspartate aminotransferase (AST) (Sigma, #MAK055), bilirubin (total and direct) (Abcam, #ab235627), and GGT activity (Abcam #ab241029) were measured according to manufacturer’s instructions.

### Bile acid (BA) profile

The bile acid (BA) pool was analyzed as previously described (64). The liver, gallbladder and small intestine were collected from 5h fasted mice and doubly homogenized in 50% methanol using a rotor-stator homogenizer, followed by a Dounce Teflon glass homogenizer.

Deuterated BA standard (20 *μ*L of 25 *μ*M d4-cholic acid) was added to 200 *μ*L of each sample, and calibrator curves were generated of each BA in charcoal-stripped tissue. To each sample/calibrator, 2 mL of ice-cold acetonitrile was added, then samples were vortexed for 1 hour at 2,000 rpm and centrifuged for 10 min at 11,000 g. Supernatants were transferred to clean glass tubes and dried down at 45C under nitrogen. Each sample/calibrator was extracted a second time in 1 mL of ice-cold acetonitrile, vortexed for 1 hour at 2,000 rpm and centrifuged for 10 min at 11,000 g. The supernatant of the second extraction was combined with the first and dried down at 45C under nitrogen. Each sample was resuspended in 200 *μ*L of 55:45 (vol/vol) methanol:water, both with 5 mM ammonium formate.

Samples were centrifuged in UltraFree MC 0.2-*μ*m centrifugal filters (Millipore) and transferred to liquid chromatography-mass spectrometry (LC-MS) vials, and 10 *μ*L were injected into ultraperformance liquid chromatography-tandem mass spectrometry vials (UPLC-MS/MS; Waters).

### Sample preparation for metal analysis

All solutions were prepared using ultrapure reagents and ultrapure water (≥18.2 MΩ cm). Ultrapure nitric acid (HNO_3_, 65-67%, 2× subboiled) was used for sample digestion and calibration standards. Ultrapure water (≥18.2 MΩ cm) from a water purification system (Hydro) was used for reagents and standard solutions.

Mouse tissues were digested using a microwave-assisted acid digestion method for total quantitative analysis of Mn, Fe, Ca, Mg, Cu, Zn, and Se. Wet tissues were weighed (±0.01 mg) into acid-cleaned 20 mL PFA vessels, followed by the addition of 2 mL HNO_3_. Quality control included bovine liver (NIST 1577C) as a reference material and blank samples containing only HNO_3_. Digestion was performed in a MARS 6 Microwave System (CEM Corp.), ramping the temperature to 200°C in four stages and held for 15 minutes. Post-digestion, samples were cooled, transferred to 15 mL metal-free polypropylene tubes (VWR, LOT 230720058AA), and diluted to 10 mL with ultrapure water. A 2 mL subsample was mixed with 40 µL internal standard (500 µg/L Ga and Y in 2% HNO_3_) and 40 µL gold (50,000 µg/L stock), then diluted to 4 mL with ultrapure water.

Mouse bile samples were weighed, then digested overnight at room temperature in the original vials with 200 µL of concentrated ultrapure HNO_3_. After digestion, samples were transferred to 15 mL metal-free polypropylene tubes, spiked with 20 µL of internal standard (500 µg/L Ga, Y in 2% HNO₃) and 20 µL of 50,000 µg/L Au stock, and diluted to 2 mL with ultrapure water. The empty original vials were then weighed to determine bile mass and dilution factor.

### Inductively Coupled Plasma-Mass Spectrometer measurement

An Agilent 8900 inductively coupled triple quadrupole plasma mass spectrometer (ICP-QQQ-MS) with an SPS 4 autosampler was used for analysis. The standard ICPMS/MS setup included a MicroMist nebulizer, double-pass spray chamber, Pt/Cu sampler and skimmer cones, and a 2.5 mm quartz plasma torch. Operating parameters: RF power 1,550 W, plasma gas flow 15.0 L/min, auxiliary gas flow 0.9 L/min, and spray chamber temperature 2°C.

External eight-point calibration was performed using matrix-matched solutions (10% HNO₃, 500 µg/L Au, 5 µg/L Ga, Y). Mg, Ca, Fe, Cu, Zn, Mn, and Se were measured in different gas modes: Cu (He mode, mass 63), Mg, Ca, Fe, Mn (NH₃ mode, mass 26, 44, 57, 66), and Se (O₂ mode, mass shift 80→96). The limit of detection (LoD) was determined as 3.33 × standard deviation of blank measurements. For digested tissues (blanks n=4), LoDs were: Mg (1.6 µg/L), Ca (59 µg/L), Mn (0.08 µg/L), Fe (5.8 µg/L), Cu (0.21 µg/L), Zn (26 µg/L), and Se (0.06 µg/L). For bile samples (blanks n=5), the LOD was 0.09 µg/L for Mn. The recovery for CRM NIST 1577c (n=7) was 96% for Mg, 97% for Ca, 82% for Mn, 89% for Fe, 82% for Cu, 89% for Zn and 98% for Se, as percentages of certified values.

### Bile flow experiments

Experiments were carried out using a surgical approach similar to ref (65). 8 week old C3H/HeJ mice (Jackson #000659) were fasted for 5-hours before being orally gavaged with water or 4 g/kg body weight fructose. 90 minutes after gavage mice were anesthetized with ketamine (100 mg/kg)/xylazine (10 mg/kg), the distal common bile duct was ligated and the gall bladder was cannulated with a 30G needle on catheter tubing (BD Intramedic PE tubing #427401) of the same length (4 in.), and tubing was diverted into pre-weighed tubes. Bile flow rate was determined by the weight of the bile, assuming a density of 1 g/ml.

### Chromatin Immunoprecipitation Sequencing (ChIP-Seq)

ChIP-sequencing experiment and analysis have been described previously (66). The data have been deposited in GEO (GSE217983).

### Primary Hepatocyte Culture

Primary mouse hepatocytes were isolated from wildtype C57BL/6J mice aged over 10 weeks using a two-step collagenase perfusion protocol (67). Mice were anesthetized using ketamine (100 mg/kg)/xylazine (10 mg/kg) IP injections. Mice were affixed and the inferior vena cava was exposed and catheterized with a 24-gauge catheter (Excel International) and infused 25 mL of EGTA-based perfusion solution followed by 25 mL of Type IV collagenase solution (Worthington #LS004188). Following cell dissociation in complete medium (Corning M199 #10060CV + 10% fetal bovine serum (Gibco #A5669801) + 1% Penicillin-Streptomycin (PenStrep, Gibco #15140148)) cells were filtered through a 70 *μ*m mesh cell strainer. Hepatocytes were isolated by Percoll (Cytvia #17089101) density gradient centrifugation steps. Viable cells were suspended in complete medium and seeded in Type I collagen (Sigma #C38671) coated plates.

For protein studies in mouse primary hepatocytes, serum-free medium (M199 + 1% PenStrep) was replaced for 16 hours overnight with or without 6 *μ*M MnCl_2_ (Fisher #M87). Serum-free medium supplemented with 10 *μ*M forskolin (Thermo Scientific Chemicals #J63292MA) and 1 *μ*M dexamethasone (Sigma #D2915) was added for 5 hours. 1 or 100 nM of insulin (Gibco #12585014) was then added for 3 minutes. Cells were washed 2x with ice cold PBS before direct cell lysis with ice cold RIPA buffer (Thermo Scientific Chemicals #J63306AK) supplemented with EDTA-free Pierce protease inhibitors (Thermo Scientific #A32965) and PhosSTOP phosphatase inhibitors (Roche). For studies with the Akt inhibitor (Figure 3A), 10 *μ*M of ipatasertib (MedChemExpress #HY-15186) or DMSO was added 1 hour prior to cell lysis.

For phosphoproteomic analysis, mouse primary hepatocytes were serum-starved for 16 hours overnight in M199 with or without 6 *μ*M MnCl_2_. Cells were treated with serum-free medium supplemented with 10 *μ*M forskolin and 1 *μ*M dexamethasone for 5 hours prior to 1 nM of insulin for 3 minutes. Cells were washed with PBS and directly lysed in a buffer containing 200 mM EPPS (pH 8.5), 8M urea and 0.1% SDS with protease and phosphatase inhibitors and genomic DNA was sheared before sample submission.

For mRNA studies in mouse primary hepatocytes, serum-free medium (M199 + 1% PenStrep) was replaced for 16 hours overnight with or without 6 *μ*M MnCl_2_. Hepatocytes were pre-treated with 1 or 100 nM of insulin (Gibco #12585014) for 1 hour prior to 10 *μ*M forskolin/1 *μ*M dexamethasone treatment in serum-free medium for 5 hours. Cells were washed 2x with ice cold PBS before direct cell lysis with TriZol (Invitrogen #15596026).

For ChREBPβ overexpression, 50 MOI of Ad.ChREBPβ or Ad.eGFP was added for 16 hours overnight in serum free media. Hepatocytes were then treated with 10 *μ*M forskolin/1 *μ*M dexamethasone treatment in serum-free medium for 5 hours. Cells were washed 2x with ice cold PBS before direct cell lysis with TriZol.

Primary human hepatocytes were obtained from AnaBios (Lot #1183, #1141, #1045) and thawed according to manufacturer’s instructions with thaw medium B (AnaBios #HEP-005) and seeded in Type I collagen coated plates in plating medium (AnaBios #HEP-003). 6-hours after plating, medium was replaced with serum-free medium (William’s E medium (Fisher #A1217601) + 0.1 *μ*M dexamethasone, 2 mM GlutaMax (Gibco #35050061) + 15 mM HEPES (Gibco #15630080)) for 16 hours overnight with or without 6 *μ*M MnCl_2_. Serum-free medium was supplemented with 10 *μ*M forskolin and 1 *μ*M dexamethasone was added for 5 hours. 1 nM of insulin (Gibco #12585014) was then added for 3 minutes. Cells were washed 2x with ice cold PBS before direct cell lysis with lysis buffer supplemented with protease and phosphatase inhibitors.

### Western blotting

Proteins were extracted from tissues by homogenizing tissues directly in RIPA buffer supplemented with protease/phosphatase inhibitors using a rotor-stator homogenizer. Protein concentration was assessed by BCA assay (Thermo Scientific #23225). An equal amount of protein was loaded into Criterion TGX polyacrylamide gels (BioRad) and transferred onto 0.2 *μ*m PVDF (BioRad) or nitrocellulose (LI-COR) membranes. Primary antibodies included: anti-phospho insulin receptor β Tyr1150 (Cell Signaling #3024), anti-insulin receptor β (Cell Signaling #3025), anti-phospho AKT Thr308 (Cell Signaling #13038), anti-phospho AKT Ser473 (Cell Signaling #9371), anti-pan AKT (Cell Signaling #4691), anti-phospho GSK3β Ser9 (Cell Signaling #5558), anti-GSK3β (Cell Signaling #12456), anti-phospho PRAS40 Thr246 (Cell Signaling #2997), anti-PRAS40 (Cell Signaling #2691), anti-phospho FoxO1 Ser256 (Cell Signaling #84192), anti-FoxO1 (Cell Signaling #2880), anti-Gapdh (Cell Signaling #2118), anti-phospho AKT Substrate (RxRxxS/T) (Cell Signaling #10001), anti-phospho PKA substrate (RRxS/T) (Cell Signaling #9624). Secondary antibody was ECL Rabbit IgG, HRP-linked whole Ab (Cytvia #NA-934) followed by Pierce ECL Western Blotting substrate (Thermo Scientific #32106). Densitometric analysis was performed using ImageJ software from the National Institutes of Health.

### RNA isolation and quantitative real-time PCR

RNA was extracted with TriZol reagent according to manufacturer’s instructions. 2 *μ*g of RNA was used to synthesize cDNA by reverse transcription using the high-capacity cDNA reverse transcription kit (Applied Biosystems #4368814). Gene expression quantification was performed by quantitative PCR with iTaq Universal SYBR Green Supermix (Biorad #1725122). Gene expression was normalized to *36b4* or *TBP* as a housekeeping gene. Primer sequences are:

*G6pc* F: CGACTCGCTATCTCCAAGTGA

*G6pc* R: GTTGAACCAGTCTCCGACCA

*Pck1* F: CTGCATAACGGTCTGGACTTC

*Pck1* R: CAGCAACTGCCCGTACTCC

*Gck* F: TGAGCCGGATGCAGAAGGA

*Gck* R: GCAACATCTTTACACTGGCCT

*Igfbp1* F: AGATCGCCGACCTCAAGAAAT

*Igfbp1* R: CTCCAGAGACCCAGGGATTTT

*Chrebpb* F: TCTGCAGATCGCGTGGAG

*Chrebpb* R: CTTGTCCCGGCATAGCAAC

*Slc30a10* F: GAGATGGGCCGTTACTCAGG

*Slc30a10* R: GCCTCCACGAAGATGGTGAA

*Slc39a8* F: CTGTCACTGAGCCTAACGGA

*Slc39a8* R: GCCGTCGATGAAATTGTGG*A*

*Slc39a14* F: ATCCAGAATCTTGGCCTCCT

*Slc39a14* R: AAGAGCTGCCTTTTCCATGA

*36b4* F: AGATGCAGCAGATCCGCAT

*36b4* R: GTTCTTGCCCATCAGCACC

### Glucose Production Assays

125,000 primary hepatocytes were seeded in a 12-well, type I collagen coated plate in complete medium (M199 + 10% FBS + 1% PenStrep). Hepatocytes were serum starved in M199 + 1% PenStrep overnight for 16 hours with or without with or without 6 *μ*M MnCl_2_. Insulin-treated wells were pre-treated with 1 or 100 nM insulin for 1 hour. Media was then aspirated, wells were washed 2x with pre-warmed PBS, then 700 *μ*L of glucose production medium was added/well with or without 10 *μ*M forskolin/1 *μ*M dexamethasone, 6 *μ*M MnCl_2_, and insulin for 7 hours. Glucose production media was glucose-free, sodium pyruvate-free, phenol red-free, glutamine-free DMEM (Gibco #A14430) supplemented with 2 mM sodium pyruvate (Gibco #11360070), 2.24 g/L sodium L-lactate (Sigma #L7022), 0.5X GlutaMax Supplement (Gibco #35050061), and 15 mM HEPES (Gibco #15630080). Glucose production media was collected in pre-chilled eppendorf tubes, spun at 3000 rpm for 5 minutes at 4C, then 400 *μ*L of supernatant was collected into new pre-chilled eppendorf tube for glucose analysis by colorimetric glucose assay (Wako AutoKit Glucose #99703001). Cells were washed with ice cold PBS and cells were directly lysed with RIPA buffer with protease/phosphatase inhibitors. Protein content was determined by BCA assay to normalize glucose production.

### RNA-Sequencing

RNA was extracted from primary hepatocyte samples using RNeasy Mini Kit (Qiagen #73404). RNA concentration and quality was determined by Qubit Bioanalyzer. RNA samples from 3 samples/treatment group were submitted to the JP Sulzberger Columbia Genome Center for library preparation (standard poly(A) pulldown for mRNA enrichment), RNA-Seq (20M depth, paired-end 75bp reads using an Element Biosciences AVITI) and processing of raw data. Differential expression analysis on counts data were done using the DESeq2 package.

### Kinase Activity Assays

In ATP consumption assays the Promega ADP-Glo Assay (#V9101) was used. The kinase buffer used was 160 mM Tris (pH 7.5), 200 *μ*M DTT, 0.4 mg/mL BSA. 5 ng of Akt1 (SignalChem #A16-14G) or PKA (Promega #V4246) was added per reaction. The reaction was incubated at room temperature for 45 minutes (Akt) or 30 minutes (PKA). The k_cat_/K_m_ (ATP) values were calculating using Michaelis-Menten equations using a non-linear fit with PRISM.

In phosphorylated GSK3β assays the Enzo Akt kinase activity kit (#ADI-EKS-400A) was used with slight modifications. The kinase buffer used was 50 mM HEPES/NaOH, 150 mM NaCl, 3mM DTT. 10 ng of Akt2 (SignalChem #A17-14G) or Akt2ΔPH (SignalChem #A17-15H) was added per reaction. The reaction was incubated at 30C for 30 minutes.

### Sample preparation and analysis for proteomics/phosphoproteomics mass spectrometry

Samples were processed by the Harvard Thermo Fisher Center for Multiplexed Proteomics core. Samples for protein analysis were prepared essentially as previously described (68, 69). Detailed methods for proteomics and phosphoproteomics analysis, peptide enrichment, and LC-MS/MS measurements are described in **Supplemental Methods.**

The phosphproteomics data was normalized by taking the ratio of the phosphosite abundance to the protein abundance. The ratios were then scaled to the mean of the protein abundance, so the average abundance of the phosphoproteomics is the same as the average abundance of the proteomics. The phosphoproteomics data was then log2 transformed. The heatmaps show the clipped Z-scores of log2 abundance of each protein in each treatment group. To determine if any phosphosite was differentially expressed between two treatment groups we performed differential expression analysis using the moderated t-test of Limma package (75). To detect additive effects of Mn and insulin, we consider phosphosites where the effect of insulin and the effect of Mn is in the same direction relative to mock, and whose p-value indicates significance of additivity using the Treat method (37) from Limma. If both insulin and Mn are not in the same direction, then we assign the additivity test statistics for that phosphosite to be NA. Motif sequence logos were created using PhosphoSitePlus (76).

For kinase and phosphatase pathway analysis, we performed gene set enrichment analysis using the gene sets of the RegPhos and PhosphositePlus databases (76, 77) for kinase substrates and the Depod database (78) for phosphatase substrates. Positive/negative Normalized Enrichment Scores (NES) indicates overall up/down-regulation of phosphosites for kinase substrates, but opposite regulation of phosphosites for phosphatase substrates.

## Statistical Analysis

Data are presented as mean ± SEM. Statistical analysis was by Student’s t-test, one-way ANOVA or two-way ANOVA followed by a Tukey’s post hoc multiple comparison test. p<0.05 is considered statistically significant. ns=not significant, *p<0.05, **p<0.01, ***p<0.001, ****p<0.0001.

## Study Approval

All experiments were approved by the Institutional Animal Care and Use Committees of Columbia University Medical Center and Duke University.

## DATA AVAILABILITY

Requests for further information and resources should be directed to and will be fulfilled by the lead contact, Rebecca Haeusler

This study did not generate new unique reagents.

## Data and code availability

- Raw RNA seq data is deposited at the Gene Expression Omnibus (GEO) and will be made publicly available as of the date of publication.
- The phosphoproteomic dataset generated for this study is deposited in the PRoteomics IDEntifications (PRIDE) database and will be made publicly available as of the date of publication.
- This paper does not report original code.
- Any additional information required to reanalyze the data reported in this paper is available from the lead contact upon request.

## AUTHOR CONTRIBUTIONS

JRG and RAH conceptualization; JRG, SH, TLY, CL, KS, EC, AG, IIA, MAH and RAH methodology; JRG, SH, TLY, YX, CL, NS, HY, MOK, SAH, KS, and IIA, data curation; JRG, TLY, YX, CL, HY, MOK, KS, and IIA, formal analysis; JRG, AN-A,EC, AG, MAH and RAH data interpretation; JRG, IIA and RAH visualization; JRG and RAH writing-original draft; JRG, SH, TLY, YX, CL, NS, HY, MOK, KS, AN-A, EC, AG, IIA, MAH, and RAH writing-review and editing; RAH supervision; JRG, MAH, and RAH funding acquisition.

## ACKNOWLEDGEMENTS

We are grateful to Utpal Pajvani, Domenico Accili, Yonghao Yu, and members of the Haeusler lab for helpful discussions. We thank Ronald Glabonjat at the Columbia University METALab for trace metals measurements. We thank Renu Nandakumar at the Columbia University Irving Institute for Clinical and Translational Research Biomarkers Core Laboratory for bile acid profiling. We thank the Thermo Fisher Center for Multiplexed Proteomics at Harvard Medical School and Jonathan Dreyfuss and Hui Pan from the Joslin Diabetes Center Bioinformatics and Biostatistics Core for work on phosphoproteomics analysis.

---

**Supplemental Figure 1.**
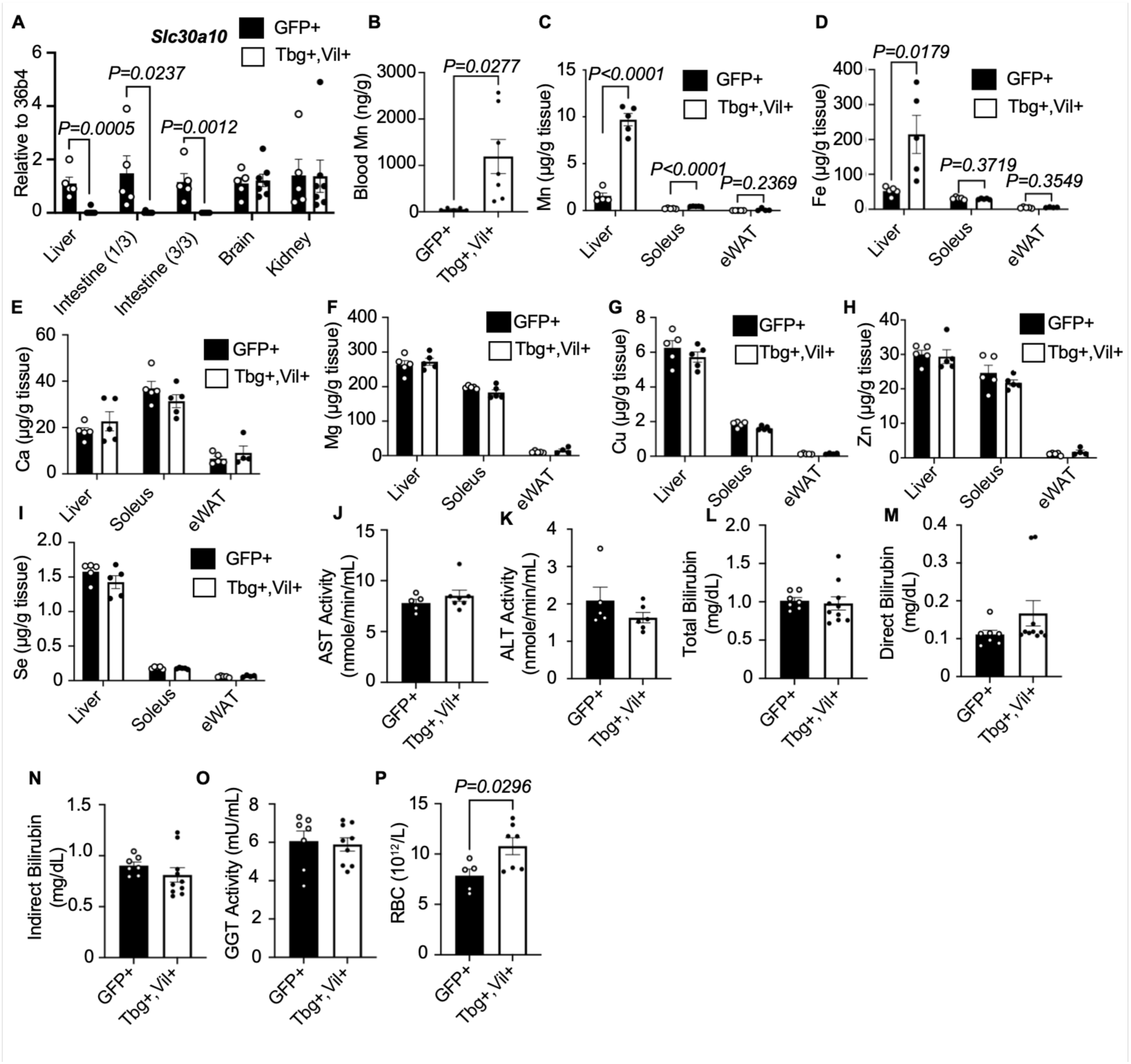
Gene expression, tissue metal levels, and liver function tests in Slc30a10^Tbg,Vil^ mice. (A) Relative gene expression of *Slc30a10* in liver, proximal (1/3) intestine, distal (3/3) intestine, brain and kidney. (B) Blood Mn levels, (C-I) liver, soleus and epididymal white adipose tissue (eWAT) metal levels from Slc30a10^Tbg,Vil^ mice, (n=4-7 males/group). Liver Mn concentration is the same as the data plotted in Figure 1A, replotted here for comparison. (J) plasma aspartate aminotransferase (AST) and (K) alanine aminotransferase (ALT) activity, (L) total bilirubin, (M) direct bilirubin, (N) indirect bilirubin, (O) GGT activity and (P) red blood cell counts from mice fed a chow diet, n=5-6 males/group. Data are presented as mean ± SEM. Student’s t-test for all panels was used.

**Supplemental Figure 2.**
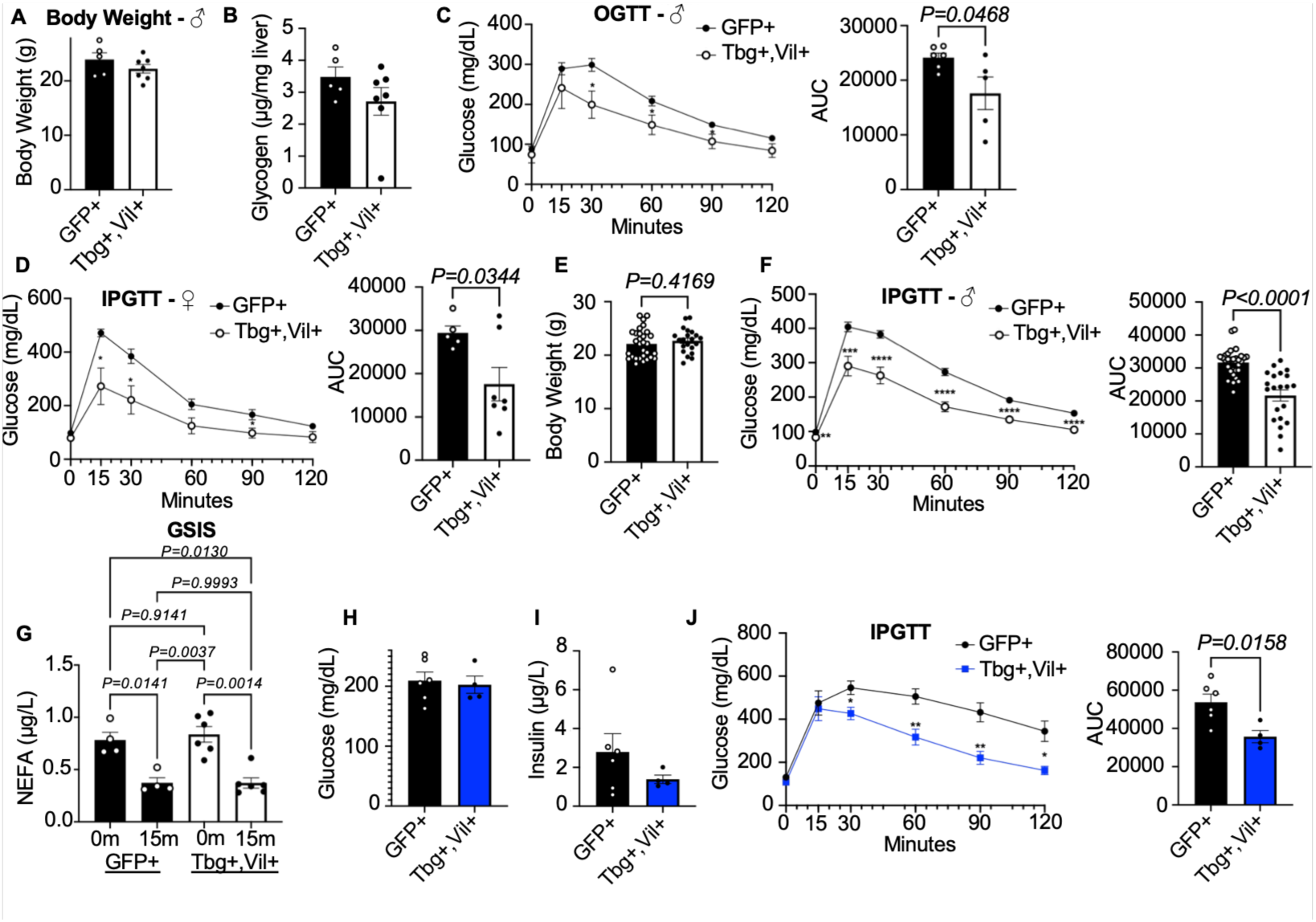
Extended metabolic parameters in Slc30a10^Tbg,Vil^ mice. (A) Body weight, (B) fasting liver glycogen, (C) oral glucose tolerance test (OGTT) from mice fed a chow diet, n=5-6 males/group. (D) Intraperitoneal glucose tolerance test (IPGTT) in mice fed a chow diet, n=5-6 females/group. (E) Body weight and (F) IPGTT from five pooled independent cohorts, comparing controls to only the Slc30a10^Tbg,Vil^ mice that had body weights within the range of the body weights of control mice. n=21-30 males/group. (G) Plasma non-esterified fatty acids during glucose-stimulated insulin secretion (GSIS) tests in mice fed a chow diet, n=5-6 females/group. (H) 5-hour fasting glucose, (I) plasma insulin, (J) IPGTT in mice fed a high-fat, high-sucrose diet for 14 weeks, n=4-6 males/group. Data are presented as mean ± SEM. Student’s t-test for all panels except (G) where two-way ANOVA was used. *p<0.05, **p<0.01, ***p<0.001, ****p<0.0001.

**Supplemental Figure 3.**
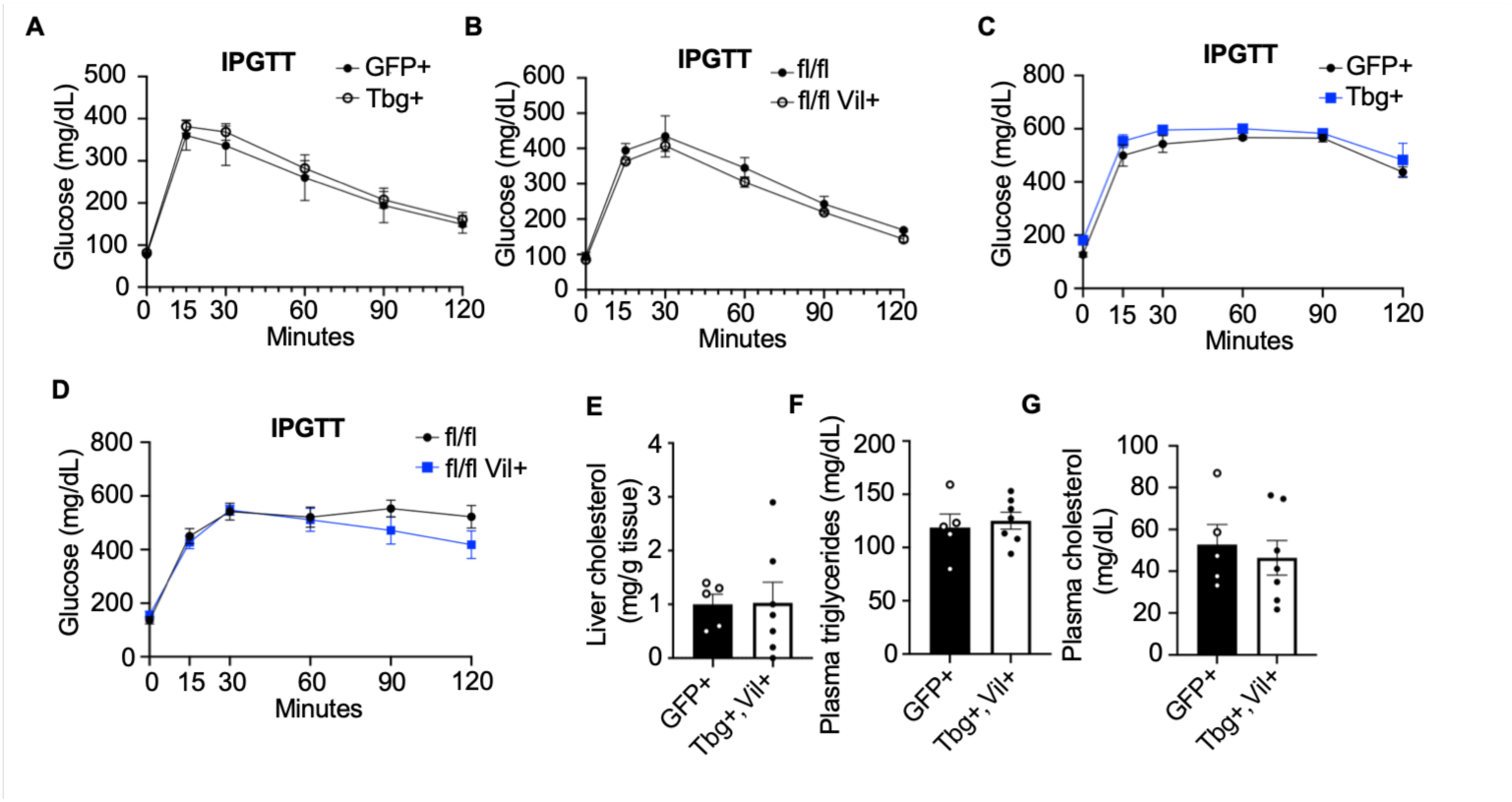
Metabolic parameters in Slc30a10^Tbg^ and Slc30a10^Vil^ mice. IPGTT in (A) Slc30a10^Tbg^ or (B) Slc30a10^Vil^ mice compared to controls fed a chow diet or a high fat, high sucrose diet for 14 weeks (C-D) n=3-5/ males/group. (E) Liver cholesterol, (F) plasma triglycerides, (G) plasma cholesterol in mice fed a chow diet, n=5-7 males/group. AUC = area under the curve. Data are presented as mean ± SEM. Student’s t-test for all panels.

**Supplemental Figure 4.**
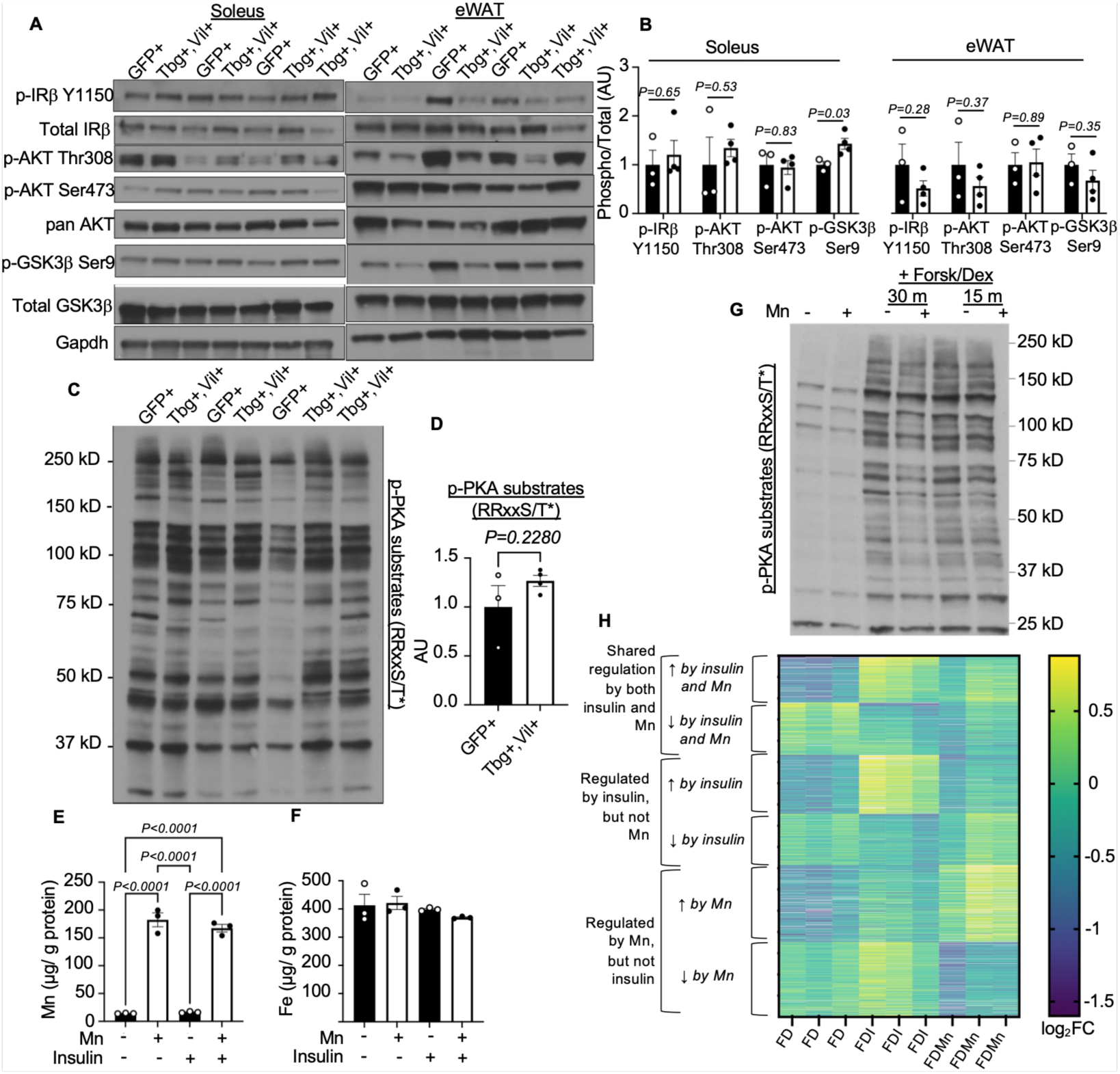
Increased Akt signaling in liver and primary hepatocytes, but not soleus or eWAT from insulin injected Slc30a10^Tbg,Vil^ mice. (A) Immunoblot analysis of the insulin signaling pathway in soleus and epididymal white adipose tissue (eWAT) with (B) densiometric quantifications from insulin-injected Slc30a10^Tbg,Vil^ mice. (C-D) Immunoblot of phosphorylated PKA substrates from liver of insulin-injected Slc30a10^Tbg,Vil^ mice with densitometric quantification. Cellular (E) Mn and (F) iron (Fe) content in primary hepatocytes treated with Mn (6 μM) or insulin (1 nM). (G) Immunoblot analysis of phosphorylated PKA substrates in primary hepatocytes treated with or without forskolin/dexamethasone (Forsk/Dex) and Mn. (H) Differentially expressed genes in RNA-sequencing of primary hepatocytes treated with forskolin/dexamethasone (FD) plus insulin (FDI) or Mn-treated (FDMn). Data included in the heatmap fit the parameters of P_adj_<0.05, AND log2FC>|0.5| for FDI or FDMn or both, compared to FD. For panels B, D, E, and F, Data are presented as mean ± SEM. Student’s t-tests were used.

**Supplemental Figure 5.**
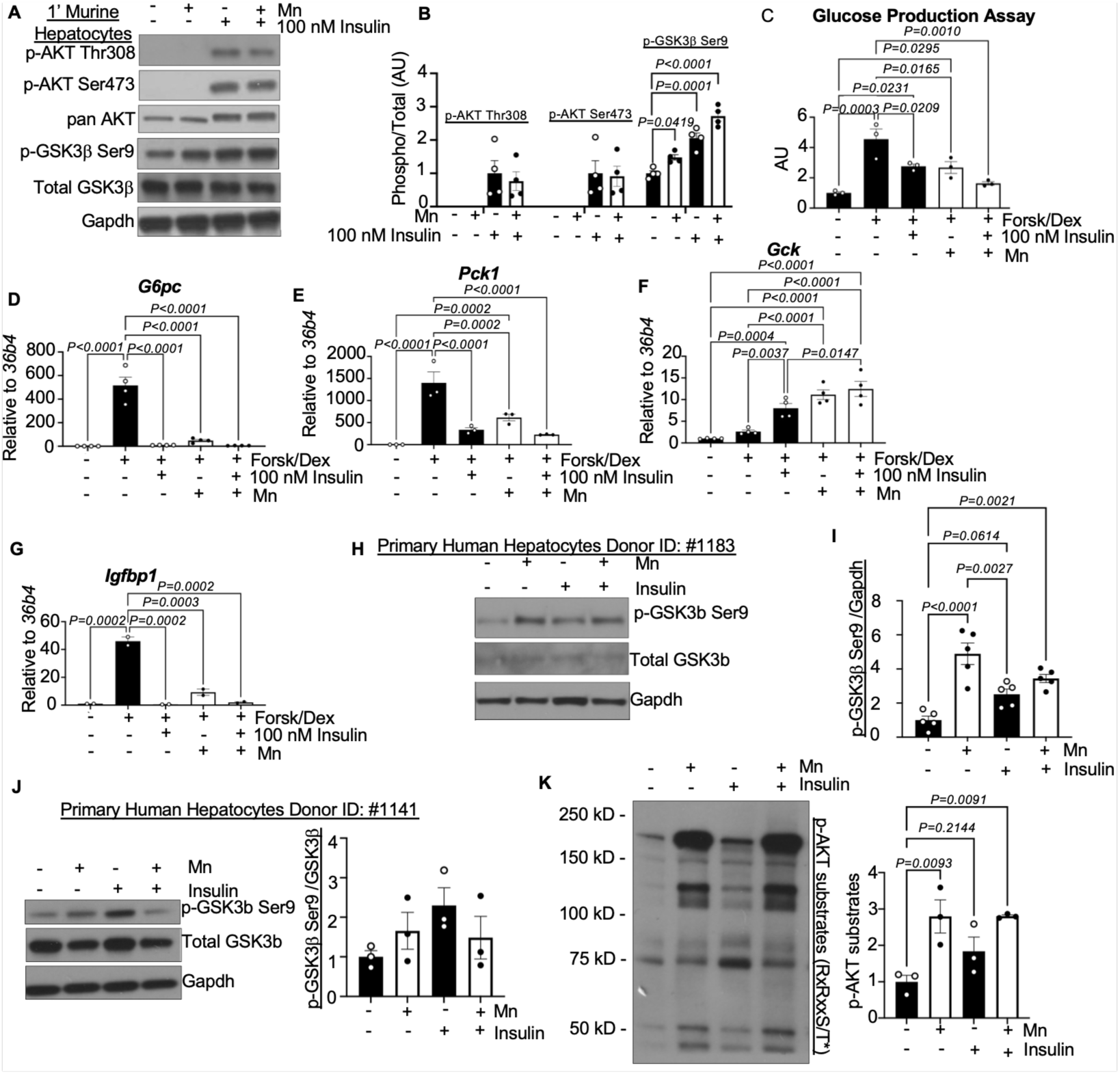
Increased Akt signaling in Mn-treated primary mouse hepatocytes. (A) Representative immunoblot analysis of the insulin signaling pathway in primary hepatocytes treated with Mn (6 μM) or insulin (100 nM). (B) Relative densitometric quantification of (A). (C) Relative glucose production and (D-G) gene expression from primary hepatocytes treated with forskolin/dexamethasone (Forsk/Dex), insulin or Mn. (H-K) Representative immunoblots from primary human hepatocytes treated with Mn (6 μM) or insulin (1 nM) with relative densitometric quantifications. Each data point represents a technical replicate from the same donor. Data are presented as mean ± SEM. One-way ANOVAs were used.

**Supplemental Figure 6.**
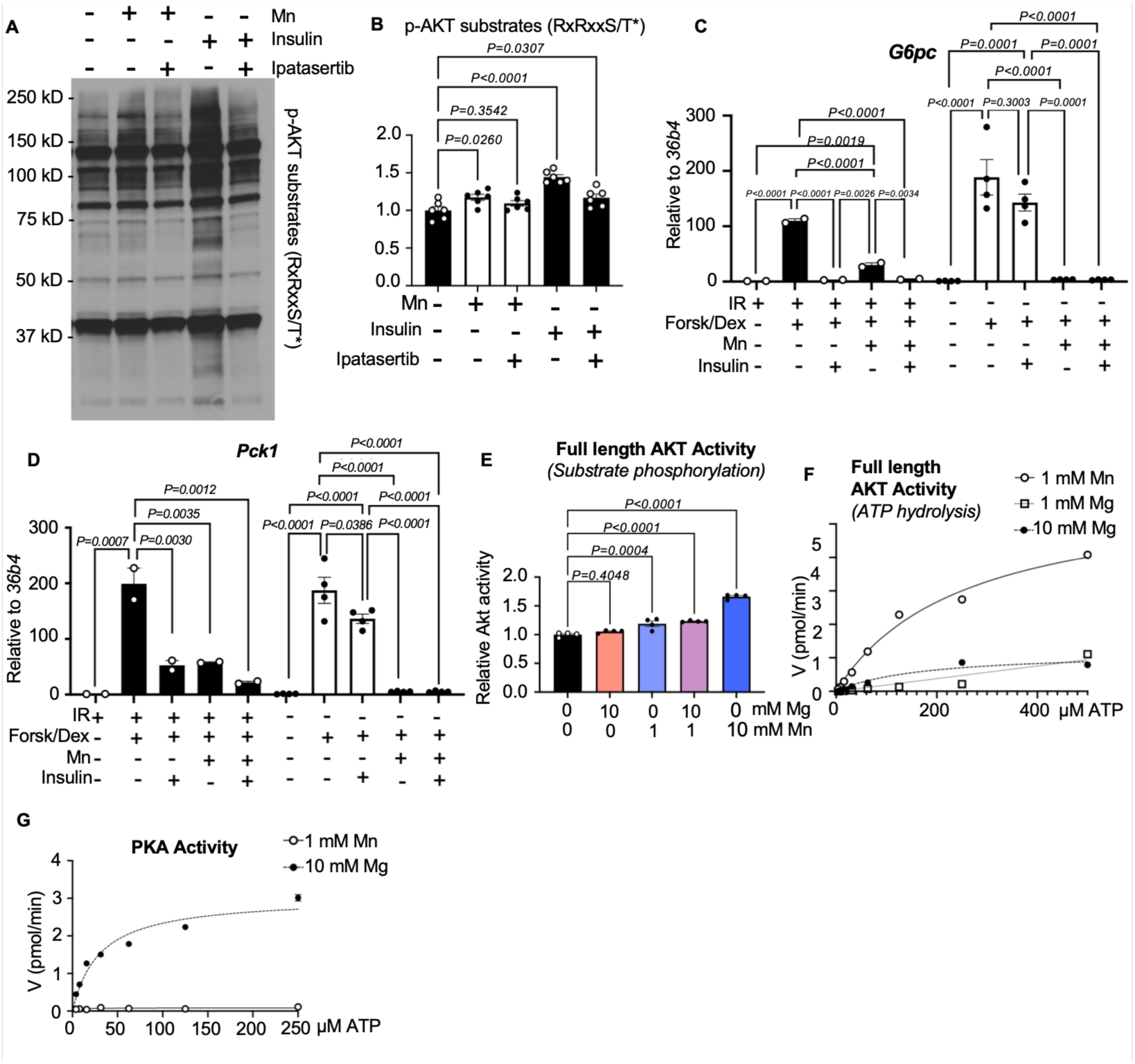
Mn acts on Akt to increase its activity. (A) Representative immunoblot analysis of phosphorylated Akt substrates from primary hepatocytes treated with insulin (1 nM), Mn (6 μM), or ipatasertib (10 μM) and (B) densitometric quantification. (C-D) Gene expression from primary hepatocytes derived from control or LIRKO mice treated with forskolin/dexamethasone (Forsk/Dex), insulin or Mn. (E) In vitro phosphorylation of Gsk3β peptide and (F) consumption of ATP by purified full-length Akt. (G) In vitro PKA activity. Data are presented as mean ± SEM. One-way ANOVAs were used.

**Supplemental Figure 7.**
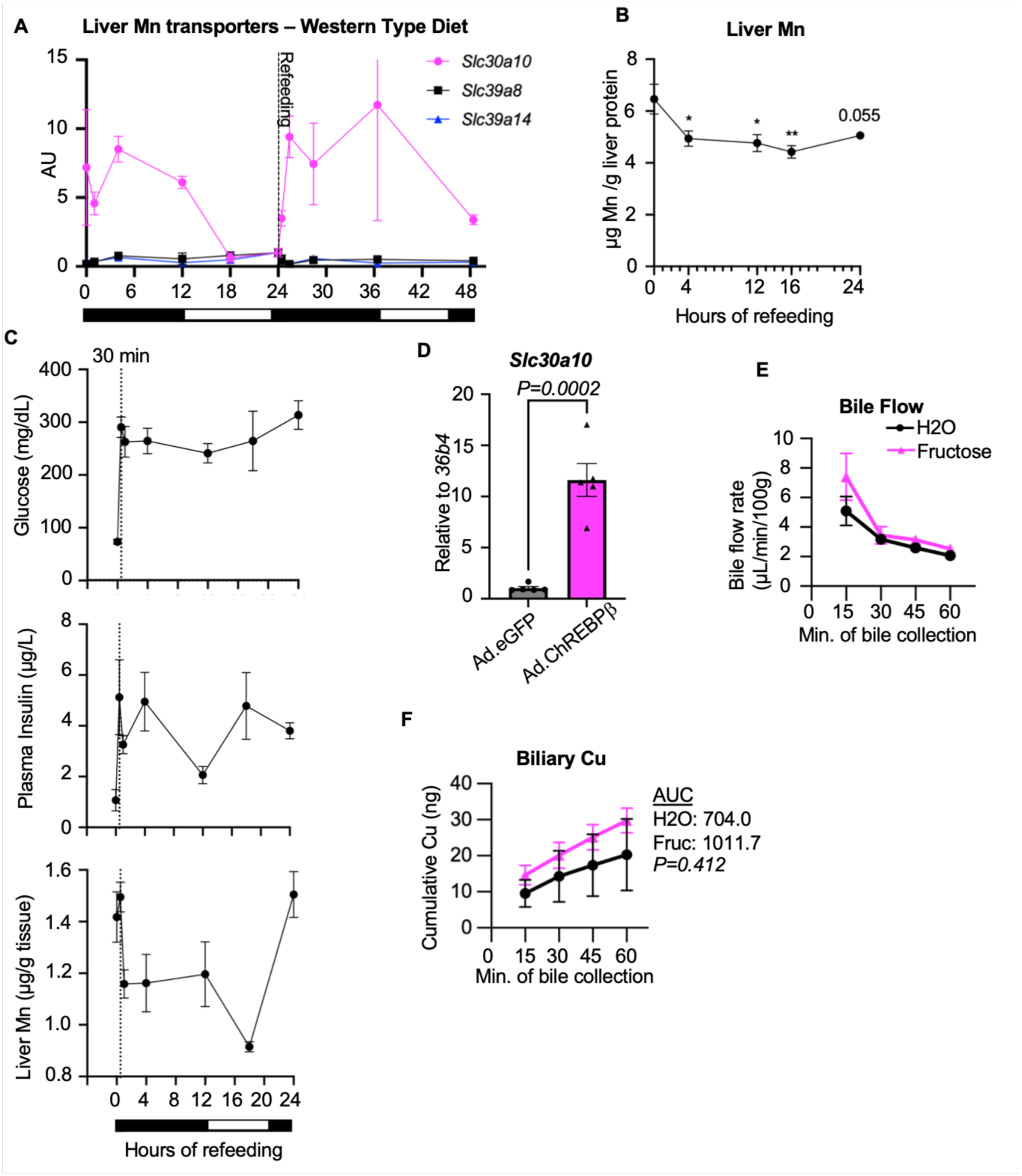
Slc30a10 is nutritionally regulated during the fasting/refeeding transition. (A) Relative gene expression from liver tissue of wild type C57BL/6J mice fed a western-type diet for one week prior to being fasted for 24 hours followed by refeeding with the same diet, n=3-5 males/group. (B) Liver Mn concentrations expressed per gram of protein during indicated refed times with chow diet after a 24 hour fast in wild type C57BL/6J mice, n=5 males/group. An independent experiment measuring (C) plasma glucose, plasma insulin and liver Mn concentrations during indicated refed times with chow diet after a 24 hour fast in wild type C57BL/6J mice, n=5 males/group. (D) Gene expression from primary mouse hepatocytes treated with control adenovirus expressing eGFP (Ad.eGFP) or 3xFlag-ChREBPβ (Ad.ChREBPβ). (E) Bile flow rates and (F) biliary copper (Cu) content at indicated time points of wildtype C3H/HeJ mice gavaged with fructose or water, n=3-4 males/group. AUC = area under the curve. Data are presented as mean ± SEM. *p<0.05, **p<0.01 by Student’s t-tests.

**Supplemental Table 1.**
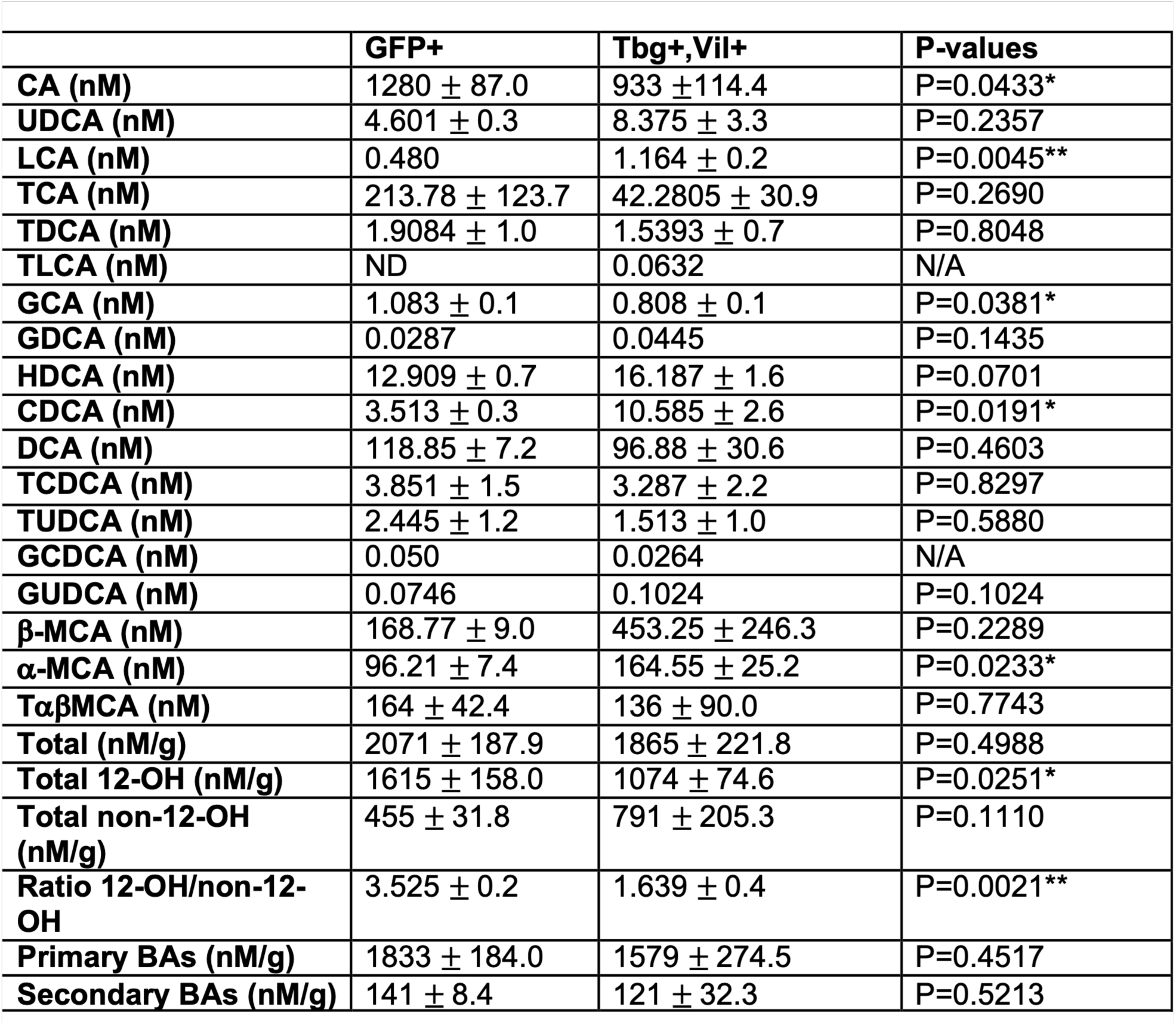
Composition of the total bile acid (BA) pool in Slc30a10^Tbg, Vil^ or control (GFP+) mice. ND = not detected, N/A = not applicable. All data are presented as mean ± SEM. *p<0.05, **p<0.01, by Student’s t-tests.

**Supplemental Table 2.** Gene list in heatmap of Supplemental Figure 4H.

**Supplemental Table 3.** RNA-seq pathway analysis using DAVID Bioinformatics tool.

**Supplemental Table 4.** Differential expression analysis of phosphorylated peptides.

**Supplemental Table 5.** Motif analyses from phosphoproteomics. For Akt recognition motifs, the motifs are mutually exclusive - a given phosphosite is listed as only one of the Akt motifs. For non-Akt recognition motifs, a phosphosite may be listed as more than one motif, as some kinases have overlapping recognition motifs. The most common overlap is MAPK motifs with CDKs or PKC.

|  | Induced by insulin<br>p<0.05<br>(300) | Significantly Additive<br>[induced by insulin<br>p<0.05 and induced<br>by Mn p<0.05 and<br>Mn+ins>Mn and<br>Mn+ins>[ins] and<br>additive effect size is<br>significant p<0.05<br>(70) | Additive<br>[induced by insulin<br>p<0.05 and induced<br>by Mn p<0.05 and<br>Mn+ins>Mn and<br>Mn+ins>[ins]<br>(79) | Induced by insulin<br>p<0.05 and induced<br>by Mn p<0.05 but<br>not additive<br>(52) | Induced by insulin<br>p<0.05 but reduced<br>by Mn p<0.05<br>(25) | Induced by insulin<br>p<0.05 but<br>unaffected by Mn<br>(74) |
| --- | --- | --- | --- | --- | --- | --- |
| **AKT** | **100 (30.30%)** | **37 (48.05%)** | **29 (32.58%)** | **6 (11.11%)** | **7 (25.93%)** | **21 (25.30%)** |
| (R-x-R-x-x-S/T*-ϕ) | 29 ( 8.79%) | 12(15.58%) | 5 ( 5.62%) | 1 (1.85%) | 3 (11.11%) | 8 ( 9.64%) |
| (R-x-R-x-x-S/T*) | 17 ( 5.15%) | 8 (10.39%) | 5 ( 5.62%) | 2 (3.70%) | 0 ( 0.00%) | 2 ( 2.41%) |
| (R-x-x-S/T*-ϕ) | 35 (10.61%) | 12 (15.58%) | 9 (10.11%) | 2 (3.70%) | 1 ( 3.70%) | 11 (13.25%) |
| (R-x-x-S/T*) | 19 ( 5.76%) | 5 ( 6.49%) | 10 (11.24%) | 1 (1.85%) | 3 (11.11%) | 0 ( 0.00%) |
| **MAPKs**<br>(P/ϕ-x-S/T*-P) | **74 (22.42%)** | **13 (16.88%)** | **17 (19.10%)** | **10 (18.52%)** | **6 (22.22%)** | **28 (33.73%)** |
| **CDKs**<br>(S/T*P-x-K/R) | **17 ( 5.15%)** | **4 ( 5.19%)** | **2 ( 2.25%)** | **3 ( 5.56%)** | **0 ( 0.00%)** | **8 ( 9.64%)** |
| **PKC**<br>(S/T*-x-R/K) | **19 ( 5.76%)** | **5 ( 6.49%)** | **5 ( 5.62%)** | **1 ( 1.85%)** | **4 (14.81%)** | **4 ( 4.82%)** |
| **CK1**<br>(D/E-D/E-D/E-x-x-S/T*-ϕ) | **0 ( 0.00%)** | **0 ( 0.00%)** | **0 ( 0.00%)** | **0 ( 0.00%)** | **0 ( 0.00%)** | **0 ( 0.00%)** |
| **CK2**<br>(S/T*-D/E-x-D/E) | **6 ( 1.82%)** | **0 ( 0.00%)** | **3 ( 3.37%)** | **2 ( 3.70%)** | **0 ( 0.00%)** | **1 ( 1.20%)** |
| **AMPK**<br>(ϕ-x-R-x-x-S*-x-x-I/L) | **5 ( 1.52%)** | **1 ( 1.30%)** | **4 ( 4.49%)** | **0 ( 0.00%)** | **0 ( 0.00%)** | **0 ( 0.00%)** |
| **PKA**<br>(R-R/K-S/T*-ϕ) | **2 ( 0.61%)** | **0 ( 0.00%)** | **0 ( 0.00%)** | **0 ( 0.00%)** | **0 ( 0.00%)** | **2 ( 2.41%)** |
| **PKG**<br>(R/K-R/K-R/K-x-S/T*) | **0 ( 0.00%)** | **0 ( 0.00%)** | **0 ( 0.00%)** | **0 ( 0.00%)** | **0 ( 0.00%)** | **0 ( 0.00%)** |
| **PhK**<br>(R-x-x-S/T*-x-ϕ-R) | **1 ( 0.30%)** | **1 ( 1.30%)** | **0 ( 0.00%)** | **0 ( 0.00%)** | **0 ( 0.00%)** | **0 ( 0.00%)** |
| **Other** | **106 (32.12%)** | **16 (20.78%)** | **29 (32.58%)** | **32 (59.26%)** | **10 (37.04%)** | **19 (22.89%)** |

**Supplemental Table 6.** Phosphosite set analysis by Kinase substrates and Phosphatase substrates.

## REFERENCES

1. CDC. National Diabetes Statistics Report [Internet]. Diabetes. 2024. https://www.cdc.gov/diabetes/php/data-research/index.html. Accessed October 2, 2025.

2. James DE, Stöckli J, Birnbaum MJ. The aetiology and molecular landscape of insulin resistance. Nat Rev Mol Cell Biol. 2021;22(11):751–771.

3. Alessi DR, et al. Characterization of a 3-phosphoinositide-dependent protein kinase which phosphorylates and activates protein kinase Balpha. Curr Biol CB. 1997;7(4):261–269.

4. Sarbassov DD, et al. Phosphorylation and Regulation of Akt/PKB by the Rictor-mTOR Complex. Science. 2005;307(5712):1098–1101.

5. James SR, et al. Specific binding of the Akt-1 protein kinase to phosphatidylinositol 3,4,5-trisphosphate without subsequent activation. Biochem J. 1996;315(Pt 3):709–713.

6. Chu N, et al. Akt Kinase Activation Mechanisms Revealed Using Protein Semisynthesis. Cell. 2018;174(4):897–907.e14.

7. Alessi DR, et al. Mechanism of activation of protein kinase B by insulin and IGF-1. EMBO J. 1996;15(23):6541–6551.

8. Manning BD, Toker A. AKT/PKB Signaling: Navigating the Network. Cell. 2017;169(3):381–405.

9. Adams JA. Kinetic and Catalytic Mechanisms of Protein Kinases. Chem Rev. 2001;101(8):2271– 2290.

10. Gurol KC, et al. Role of excretion in manganese homeostasis and neurotoxicity: a historical perspective. Am J Physiol-Gastrointest Liver Physiol. 2022;322(1):G79–G92.

11. Leyva-Illades D, et al. SLC30A10 Is a Cell Surface-Localized Manganese Efflux Transporter, and Parkinsonism-Causing Mutations Block Its Intracellular Trafficking and Efflux Activity. J Neurosci. 2014;34(42):14079–14095.

12. Taylor CA, et al. SLC30A10 manganese transporter in the brain protects against deficits in motor function and dopaminergic neurotransmission under physiological conditions. Met Integr Biometal Sci. 2023;15(4):mfad021.

13. Mercadante CJ, et al. Manganese transporter Slc30a10 controls physiological manganese excretion and toxicity. J Clin Invest. 2019;129(12):5442–5461.

14. Tuschl K, et al. Syndrome of Hepatic Cirrhosis, Dystonia, Polycythemia, and Hypermanganesemia Caused by Mutations in SLC30A10, a Manganese Transporter in Man. Am J Hum Genet. 2012;90(3):457–466.

15. Quadri M, et al. Mutations in SLC30A10 Cause Parkinsonism and Dystonia with Hypermanganesemia, Polycythemia, and Chronic Liver Disease. Am J Hum Genet. 2012;90(3):467–477.

16. Liu C, et al. Up-regulation of the manganese transporter SLC30A10 by hypoxia-inducible factors defines a homeostatic response to manganese toxicity. Proc Natl Acad Sci U S A. 2021;118(35):e2107673118.

17. Ahmad TR, et al. Bile acid composition regulates the manganese transporter Slc30a10 in intestine. J Biol Chem. 2020;295(35):12545–12558.

18. Li S, et al. Analysis of 1,25-Dihydroxyvitamin D3 Genomic Action Reveals Calcium-Regulating and Calcium-Independent Effects in Mouse Intestine and Human Enteroids. Mol Cell Biol. 2020;41(1):e00372–20.

19. Claro da Silva T, et al. Vitamin D3 transactivates the zinc and manganese transporter SLC30A10 *via* the Vitamin D receptor. J Steroid Biochem Mol Biol. 2016;163:77–87.

20. Taylor CA, et al. SLC30A10 transporter in the digestive system regulates brain manganese under basal conditions while brain SLC30A10 protects against neurotoxicity. J Biol Chem. 2019;294(6):1860–1876.

21. Prajapati M, et al. Hepatic HIF2 is a key determinant of manganese excess and polycythemia in SLC30A10 deficiency. JCI Insight;9(10):e169738.

22. Horie Y, et al. Hepatocyte-specific Pten deficiency results in steatohepatitis and hepatocellular carcinomas. J Clin Invest. 2004;113(12):1774–1783.

23. Haeusler RA, et al. Integrated Control Of Hepatic Lipogenesis Vs. Glucose Production Requires FoxO Transcription Factors. Nat Commun. 2014;5:5190.

24. Ono H, et al. Hepatic Akt Activation Induces Marked Hypoglycemia, Hepatomegaly, and Hypertriglyceridemia With Sterol Regulatory Element Binding Protein Involvement. Diabetes. 2003;52(12):2905–2913.

25. Haeusler RA, McGraw TE, Accili D. Biochemical and cellular properties of insulin receptor signalling. Nat Rev Mol Cell Biol. 2018;19(1):31–44.

26. Haeusler RA, et al. Impaired Generation Of 12-Hydroxylated Bile Acids Links Hepatic Insulin Signaling With Dyslipidemia. Cell Metab. 2012;15(1):65–74.

27. Haeusler RA, et al. Human Insulin Resistance Is Associated With Increased Plasma Levels of 12α-Hydroxylated Bile Acids. Diabetes. 2013;62(12):4184–4191.

28. Semova I, et al. Insulin prevents hypercholesterolemia by suppressing 12α-hydroxylated bile acids. Circulation. 2022;145(13):969–982.

29. Shaw AL, et al. ATP-competitive and allosteric inhibitors induce differential conformational changes at the autoinhibitory interface of Akt1. Structure. 2023;31(3):343–354.e3.

30. Lin K, et al. An ATP-Site On-Off Switch That Restricts Phosphatase Accessibility of Akt. Sci Signal. 2012;5(223):ra37–ra37.

31. Feehan R, et al. MAHOMES II: A webserver for predicting if a metal binding site is enzymatic. Protein Sci. 2023;32(4):e4626.

32. Reinhardt R, Leonard TA. A critical evaluation of protein kinase regulation by activation loop autophosphorylation. eLife. 2023;12:e88210.

33. Yang J, et al. Crystal structure of an activated Akt/Protein Kinase B ternary complex with GSK3-peptide and AMP-PNP. Nat Struct Biol. 2002;9(12):940–944.

34. Yang CS, et al. Hypothalamic AMP-Activated Protein Kinase Regulates Glucose Production. Diabetes. 2010;59(10):2435–2443.

35. Perry RJ, et al. Hepatic Acetyl CoA Links Adipose Tissue Inflammation to Hepatic Insulin Resistance and Type 2 Diabetes. Cell. 2015;160(4):745–758.

36. Li J-X, Cummins CL. Fresh insights into glucocorticoid-induced diabetes mellitus and new therapeutic directions. Nat Rev Endocrinol. 2022;18(9):540–557.

37. McCarthy DJ, Smyth GK. Testing significance relative to a fold-change threshold is a TREAT. Bioinformatics. 2009;25(6):765–771.

38. Humphrey SJ, Azimifar SB, Mann M. High-throughput phosphoproteomics reveals in vivo insulin signaling dynamics. Nat Biotechnol. 2015;33(9):990–995.

39. Kim M-S, et al. ChREBP regulates fructose-induced glucose production independently of insulin signaling. J Clin Invest. 2019;126(11):4372–4386.

40. Sargsyan A, et al. HGFAC is a ChREBP-regulated hepatokine that enhances glucose and lipid homeostasis. JCI Insight;8(1):e153740.

41. Yu C-P, et al. Positional distribution of transcription factor binding sites in the human genome. PLOS One. 2025;20(7):e0329226.

42. Yu C-P, et al. Discovering unknown human and mouse transcription factor binding sites and their characteristics from ChIP-seq data. Proc Natl Acad Sci U S A. 2021;118(20):e2026754118.

43. Koudritsky M, Domany E. Positional distribution of human transcription factor binding sites. Nucleic Acids Res. 2008;36(21):6795–6805.

44. Linder MC. Copper Homeostasis in Mammals, with Emphasis on Secretion and Excretion. A Review. Int J Mol Sci. 2020;21(14):4932.

45. Lin W, et al. Hepatic metal ion transporter ZIP8 regulates manganese homeostasis and manganese-dependent enzyme activity. J Clin Invest. 2017;127(6):2407–2417.

46. Park JH, et al. SLC39A8 Deficiency: A Disorder of Manganese Transport and Glycosylation. Am J Hum Genet. 2015;97(6):894–903.

47. Liu Q, Barker S, Knutson MD. Iron and manganese transport in mammalian systems. Biochim Biophys Acta BBA - Mol Cell Res. 2021;1868(1):118890.

48. Prajapati M, et al. Overlaps in mammalian iron and manganese homeostasis: recent advances. Biometals. 2026;39(3):891–906.

49. Gurol KC, et al. Role of excretion in manganese homeostasis and neurotoxicity: a historical perspective. Am J Physiol - Gastrointest Liver Physiol. 2022;322(1):G79–G92.

50. Rubenstein AH, Levin NW, Elliott GA. Manganese-induced hypoglycaemia. Lancet Lond Engl. 1962;2(7270):1348–1351.

51. Hassanein M, et al. Chronic Manganism: Preliminary Observations on Glucose Tolerance and Serum Proteins. Br J Ind Med. 1966;23(1):67–70.

52. Lee S-H, et al. Manganese Supplementation Protects Against Diet-Induced Diabetes in Wild Type Mice by Enhancing Insulin Secretion. Endocrinology. 2013;154(3):1029.

53. Lučić I, et al. Conformational sampling of membranes by Akt controls its activation and inactivation. Proc Natl Acad Sci U S A. 2018;115(17):E3940–E3949.

54. Knape MJ, et al. Divalent metal ions control activity and inhibition of protein kinases. Metallomics. 2017;9(11):1576–1584.

55. Zick Y, et al. Characterization of insulin-mediated phosphorylation of the insulin receptor in a cell-free system. J Biol Chem. 1983;258(1):75–80.

56. White MF, et al. Kinetic properties and sites of autophosphorylation of the partially purified insulin receptor from hepatoma cells. J Biol Chem. 1984;259(1):255–264.

57. Nicastro R, et al. Manganese is a physiologically relevant TORC1 activator in yeast and mammals. eLife. 2022;11:e80497.

58. Bryan MR, et al. Manganese acts upon Insulin/IGF receptors to phosphorylate AKT and increase glucose uptake in Huntington’s Disease cells. Mol Neurobiol. 2020;57(3):1570–1593.

59. Tang X, et al. IGF/mTORC1/S6 Signaling Is Potentiated and Prolonged by Acute Loading of Subtoxicological Manganese Ion. Biomolecules. 2023;13(8):1229.

60. Shi J-H, et al. Fructose overconsumption impairs hepatic manganese homeostasis and ammonia disposal. Nat Commun. 2023;14:7934.

61. Kim M, et al. Intestinal, but not hepatic, ChREBP is required for fructose tolerance. JCI Insight. 2017;2(24). 10.1172/jci.insight.96703.

62. White PJ, et al. The BCKDH Kinase and Phosphatase Integrate BCAA and Lipid Metabolism via Regulation of ATP-Citrate Lyase. Cell Metab. 2018;27(6):1281–1293.e7.

63. Folch J, Lees M, Stanley GHS. A SIMPLE METHOD FOR THE ISOLATION AND PURIFICATION OF TOTAL LIPIDES FROM ANIMAL TISSUES. J Biol Chem. 1957;226(1):497–509.

64. Oteng A-B, et al. Cyp2c-deficiency depletes muricholic acids and protects against high-fat diet-induced obesity in male mice but promotes liver damage. Mol Metab. 2021;53:101326.

65. Oude Elferink RP, et al. Regulation of biliary lipid secretion by mdr2 P-glycoprotein in the mouse. J Clin Invest. 1995;95(1):31–38.

66. Sargsyan A, et al. HGFAC is a ChREBP-regulated hepatokine that enhances glucose and lipid homeostasis. JCI Insight. 2023;8(1). 10.1172/jci.insight.153740.

67. Langlet F, et al. Selective Inhibition of FOXO1 Activator/Repressor Balance Modulates Hepatic Glucose Handling. Cell. 2017;171(4):824–835.e18.

68. Navarrete-Perea J, et al. SL-TMT: A Streamlined Protocol for Quantitative (Phospho)proteome Profiling using TMT-SPS-MS3. J Proteome Res. 2018;17(6):2226–2236.

69. Li J, et al. TMTpro-18plex: The Expanded and Complete Set of TMTpro Reagents for Sample Multiplexing. J Proteome Res. 2021;20(5):2964–2972.

70. Hughes CS, et al. Single-pot, solid-phase-enhanced sample preparation for proteomics experiments. Nat Protoc. 2019;14(1):68–85.

71. Schweppe DK, et al. Full-Featured, Real-Time Database Searching Platform Enables Fast and Accurate Multiplexed Quantitative Proteomics. J Proteome Res. 2020;19(5):2026–2034.

72. Schweppe DK, et al. Characterization and Optimization of Multiplexed Quantitative Analyses Using High-Field Asymmetric-Waveform Ion Mobility Mass Spectrometry. Anal Chem. 2019;91(6):4010–4016.

73. Rad R, et al. Improved Monoisotopic Mass Estimation for Deeper Proteome Coverage. J Proteome Res. 2021;20(1):591–598.

74. Gassaway BM, et al. A multi-purpose, regenerable, proteome-scale, human phosphoserine resource for phosphoproteomics. Nat Methods. 2022;19(11):1371–1375.

75. Ritchie ME, et al. limma powers differential expression analyses for RNA-sequencing and microarray studies. Nucleic Acids Res. 2015;43(7):e47.

76. Hornbeck PV, et al. PhosphoSitePlus, 2014: mutations, PTMs and recalibrations. Nucleic Acids Res. 2015;43(Database issue):D512–D520.

77. Lee T-Y, et al. RegPhos: a system to explore the protein kinase–substrate phosphorylation network in humans. Nucleic Acids Res. 2011;39(Database issue):D777–D787.

78. Duan G, Li X, Köhn M. The human DEPhOsphorylation database DEPOD: a 2015 update. Nucleic Acids Res. 2015;43(Database issue):D531–D535.

