## Supplemental Methods for "Manganese availability determines insulin sensitivity by enhancing Akt activity"

Sample preparation for proteomics/phosphoproteomics mass spectrometry.

Proteomes were extracted using a buffer containing 200 mM EPPS pH 8.5, 8M urea, 0.1% SDS and protease inhibitors. Following lysis, 150 µg of each proteome was reduced with 5 mM TCEP (Thermo Scientific #77720). Cysteine residues were alkylated using 10 mM iodoacetimide (MP Biochemicals #100351) for 20 minutes at room temperature in the dark. Excess iodoacetimide was quenched with 10 mM DTT (Alfa Aesar #A15797). A buffer exchange was carried out using a modified SP3 protocol(70). Briefly, ~2000 µg of Cytiva SpeedBead Magnetic Carboxylate Modified Particles (Cytvia #65152105050250 and 4515210505250), mixed at a 1:1 ratio, were added to each sample. 100% ethanol was added to each sample to achieve a final ethanol concentration of at least 50%. Samples were incubated with gentle shaking for 15 minutes. Samples were washed three times with 80% ethanol. Protein was eluted from SP3 beads using 200 mM EPPS pH 8.5 containing Lys-C (Wako #129-02541). Samples were digested overnight at room temperature with vigorous shaking. The next morning trypsin (ThermoFisher Scientific) was added to each sample and further incubated for 6 hours at 37º C. Acetonitrile was added to each sample to achieve a final concentration of ~33%. Each sample was labelled, in the presence of SP3 beads, with ~300 µg of TMTPro reagents (ThermoFisher Scientific #A44520 (1-16); #A52045 (17-18)). Following confirmation of satisfactory labelling (>97%), excess TMT was quenched by addition of hydroxylamine to a final concentration of 0.3%. The full volume from each sample was pooled and acetonitrile was removed by vacuum centrifugation for 1 hour. The pooled sample was acidified and peptides were de-salted using a Sep-Pak 50mg tC18 cartridge (Waters #WAT0954925). Peptides were eluted in 70% acetonitrile, 1% formic acid and dried by vacuum centrifugation.

Phosphopeptide enrichment. A phosphopeptide enrichment was performed using a High-Select Fe-NTA Phosphopeptide Enrichment Kit (ThermoFisher Scientific #A32992). Dried phosphopeptides were de-salted by Stage-Tip and re-dissolved in 5% formic acid/5% acetonitrile for LC-MS/MS. The flow through from the phosphopeptide enrichment was used for total proteome profiling.

Basic pH reversed-phase separation (BPRP). TMT labeled peptides were solubilized in 5% acetonitrile/10 mM ammonium bicarbonate, pH 8.0 and ~300 µg of TMT labeled peptides were separated by an Agilent 300 Extend C18 column (3.5 μm particles, 4.6 mm ID and 250 mm in length). An Agilent 1260 binary pump coupled with a photodiode array (PDA) detector (Thermo Scientific) was used to separate the peptides. A 45 minute linear gradient from 10% to 40% acetonitrile in 10 mM ammonium bicarbonate pH 8.0 (flow rate of 0.6 mL/min) separated the peptide mixtures into a total of 96 fractions (36 seconds). A total of 96 Fractions were consolidated into 24 samples in a checkerboard fashion and vacuum dried to completion. Each sample was desalted via Stage Tips and re-dissolved in 5% formic acid/ 5% acetonitrile for LC-MS3 analysis.

Liquid chromatography separation and tandem mass spectrometry (LC-MS3) for proteomics. Proteome data was collected on an Orbitrap Eclipse mass spectrometer (ThermoFisher Scientific) coupled to a Proxeon EASY-nLC 1000 LC pump (ThermoFisher Scientific). Fractionated peptides were separated using a 120 min gradient at 400 nL/min on a 35 cm column (i.d. 100 μm, Accucore, 2.6 μm, 150 Å) packed in-house. A FAIMS device enabled during data acquisition with compensation voltages set as −40, −60, and −80 V. MS1 data were collected in the Orbitrap (60,000 resolution; maximum injection time 50 ms; AGC 4 × 10^5^). Charge states between 2 and 5 were required for MS2 analysis in the ion trap, and a 120 second dynamic exclusion window was used. Top 10 MS2 scans were performed in the ion trap with CID fragmentation (isolation window 0.5 Da; Turbo; NCE 35%; maximum injection time 35 ms; AGC 1 × 10^4^). Real-time search was used to trigger MS3 scans for quantification (71). MS3 scans were collected in the Orbitrap using a resolution of 50,000, NCE of 55%, maximum injection time of 250 ms, and AGC of 1.25 × 10^5^. The close out was set at two peptides per protein per fraction (71).

Liquid chromatography separation and tandem mass spectrometry (LC-MS2) for phosphoproteomics. Phosphorylation data were collected on an Orbitrap Eclipse mass spectrometer (ThermoFisher Scientific) coupled to a Proxeon EASY-nLC 1000 LC pump (ThermoFisher Scientific). Fractionated peptides were separated using a 120 min gradient at 525 nL/min on a 35 cm column (i.d. 100 μm, Accucore, 2.6 μm, 150 Å) packed in-house. A FAIMS device enabled during data acquisition with compensation voltages set as −40, −60, and −80 V for the first shot and −45 and −65 V for the second shot(72). MS1 data were collected in the Orbitrap (120,000 resolution; maximum injection time set to auto; AGC 4 × 10^5^). Charge states between 2 and 5 were required for MS2 analysis, and a 120 second dynamic exclusion window was used. Cycle time was set at 1 second. MS2 scans were performed in the Orbitrap with HCD fragmentation (isolation window 0.5 Da; 50,000 resolution; NCE 36%; maximum injection time 300 ms; AGC 2 × 10^5^).

Phosphoproteomics data analysis. Raw files were converted to mzXML, and monoisotopic peaks were re-assigned using Monocle (73). Searches were performed using the Comet search algorithm against a mouse database downloaded from Uniprot in May 2021. We used a 50 ppm precursor ion tolerance, 1.0005 fragment ion tolerance, and 0.4 fragment bin offset for MS2 scans collected in the ion trap, and 0.02 fragment ion tolerance; 0.00 fragment bin offset for MS2 scans collected in the Orbitrap. TMTpro on lysine residues and peptide N-termini (+304.2071 Da) and carbamidomethylation of cysteine residues (+57.0215 Da) were set as static modifications, while oxidation of methionine residues (+15.9949 Da) was set as a variable modification. For phosphorylated peptide analysis, +79.9663 Da was set as a variable modification on serine, threonine, and tyrosine residues.

Each run was filtered separately to 1% False Discovery Rate (FDR) on the peptide-spectrum match (PSM) level. Then proteins were filtered to the target 1% FDR level across the entire combined data set. Phosphorylation site localization was determined using the AScorePro algorithm (74). For reporter ion quantification, a 0.003 Da window around the theoretical m/z of each reporter ion was scanned, and the most intense m/z was used. Reporter ion intensities were adjusted to correct for isotopic impurities of the different TMTpro reagents according to manufacturer specifications. Peptides were filtered to include only those with a summed signal-to-noise (SN) ≥ 180 across all TMT channels. An extra filter of an isolation specificity (“isolation purity”) of at least 0.5 in the MS1 isolation window was applied for the phosphorylated peptide analysis. For each protein or phosphorylation site, the filtered peptide TMTpro SN values were summed to generate protein or phosphorylation site quantification values. The signal-to-noise (S/N) measurements of peptides assigned to each protein were summed (for a given protein).
